# DNA replication initiation causally sets the added mass for cell division in *Escherichia coli*

**DOI:** 10.64898/2026.02.21.703636

**Authors:** Hai Zheng, Qianandong Cao, Zheng Zhang, Yang Bai, Dengjin Li, Wenqi Huang, Shuqiang Huang, Xiongfei Fu, Arieh Zaritsky, Chenli Liu

## Abstract

Faithful genome inheritance requires cell division to be precisely coordinated with DNA replication. The classical Cooper-Helmstetter paradigm posited that division occurs at a roughly constant time after replication initiation (initiation-to-division “ID-timer” model). This has since been challenged by conflicting “adder” phenomena, leaving the primary signal for division a fundamental enigma. Here, we resolve these discrepancies by establishing that cell division in *Escherichia coli* obeys a universal “ID-adder” rule: addition of a defined size increment from replication initiation. This rule is invariant across the physiological range of nutrition-affected growth rates, whereas other versions of the adder phenomenon are emergent properties shaped by inter-cycle correlations. Through population-level perturbations of initiation mass, division machinery, and replication termination, we demonstrate that replication initiation directly and causally sets the division schedule. This principle operates independently of replication termination and division protein levels, providing a robust mechanism for coordinating the genome duplication with the cell division cycle.

---

Maintaining genomic integrity requires the temporal and spatial coordination between DNA replication and cell division (*1-4*). This universal challenge for proliferating cells is elegantly simplified in bacteria, making them a powerful tool for uncovering fundamental principles of cell cycle control (*5-8*). For decades, our understanding of the bacterial cell cycle has been guided by the classic investigations of Cooper, Helmstetter, and Donachie (*1, 6, 9-11*). Their results established that DNA replication initiates at a roughly-constant cell mass per origin (*M*_*i*_), and that division follows a relatively invariant period, yielding the replication-Initiation-to-Division (“ID timer”) paradigm (*1, 12, 13*).

The advent of single-cell microscopy, however, revealed phenomena inconsistent with a simple timer model. The discovery of “adder” phenomena, where cells add a constant mass between key events, suggested a mass-based control mechanism (*1, 14-17*), but the field has fragmented around three principal adder correlations, division-to-division (DD) (*14, 17*), initiation-to-initiation (II) (*6, 18*), and initiation-to-division (ID) (*19*). The concurrent existence of these correlations fragmented the field surrounding a fundamental, yet-unanswered question: What is the primary, causal event that triggers cell division: completion of the previous division cycle or the initiation or the termination of DNA replication?

Multiple models have been proposed to reconcile these observations (*15, 18, 20-24*). Notably, the “double-adder” model posits two independent adders, for the initiation and division cycles (*19*). While it captures certain correlations, it hypothesizes, but does not experimentally establish, a causal hierarchy between initiation and division. Moreover, it fails to explain the pronounced growth-rate dependence of the DD adder, and leaves the origin of these correlations themselves unaddressed. Therefore, a unifying and causal principle that explains the hierarchy, the growth-rate dependence, and the robustness of these adder phenomena has remained elusive.

Here, we suggest that the critical link lies between initiation and division of the same cycle. We propose that the ID adder constitutes the core, invariant principle, and the apparent inconsistencies in other adder correlations arise from the inherent correlations of overlapping cell cycles. To test this hypothesis and establish causality, we employed a comprehensive strategy combining extensive single-cell analyses across diverse growth conditions and targeted genetic perturbations in *Escherichia coli*. This approach allowed us to determine the hierarchy of cell cycle control mechanisms, reconcile adder variability, and test whether the ID adder principle works independently of key checkpoints like replication termination and the core division machinery.

## The initiation-to-division adder is robust across growth rates

The discovery of adder phenomena challenged the classic timer model but also introduced complexity: multiple, potentially conflicting adder models (DD, II, and ID) have been proposed, and their reported behavior varies with growth conditions (*1, 16*). This inconsistency raises a fundamental question: is any single adder principle truly universal? To resolve this and identify a core mechanism, we first sought to systematically quantify the robustness of the DD, II, and ID adders across the physiological range of growth rates in *E. coli*.

We engineered a strain where the SeqA protein, which binds newly replicated DNA, was fused to the red fluorescent protein mScarlet-I (*25*). This allowed the timing of replication-initiation to be visualized by the appearance of fluorescent SeqA foci (*22, 26, 27*). A comprehensive single-cell assay, integrating microfluidics, time-lapse microscopy, and custom image analysis, was used to quantify growth, replication initiation, and division dynamics across 10 nutrient conditions (fig. S1A, see Supplementary Materials). The resulting dataset encompasses steady-state growth rates from 0.25 to 1.60 h^−1^ (doubling times ranging from 169 min to 26 min) (Fig. 1A). Validation experiments confirmed that cells in this system maintained physiological steady-state growth, with key parameters consistent with population-scale measurements (fig. S1B-E).

**Figure 1.**
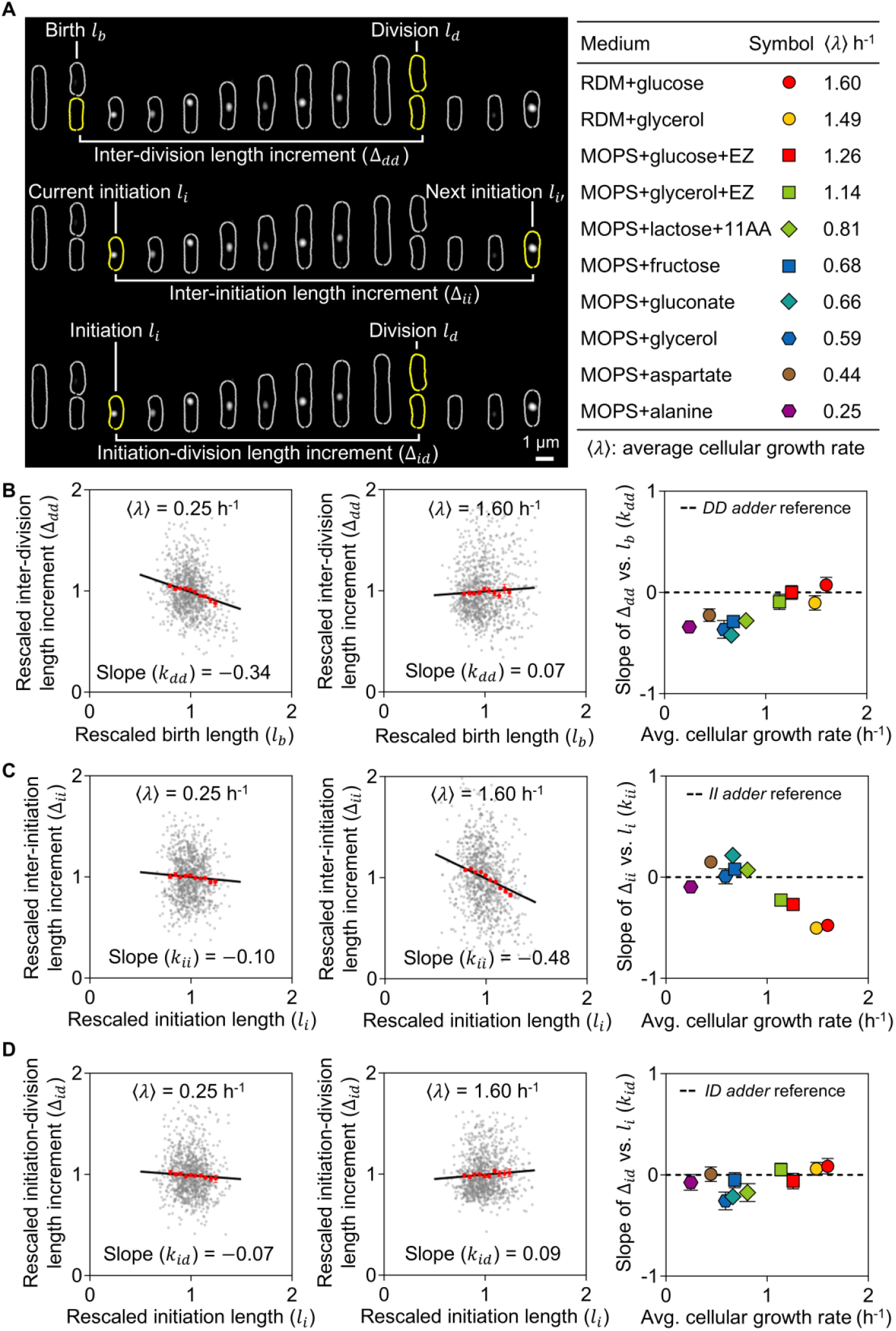
The initiation-to-division adder is robust across growth rates, in contrast to the division-to-division and initiation-to-initiation adders. (**A**) Schematic of measured parameters: birth size (*l*_*b*_), division size (*l*_*d*_), size at replication initiation (*l*_*i*_), and added sizes for division-to-division (Δ_*dd*_), initiation-to-initiation (Δ_*ii*_), and initiation-to-division (Δ_*id*_) intervals. Table summarizes growth media and average growth rates (⟨*λ*⟩). (**B**-**D**) Added size (Δ) versus initial size for (B) division-to-division (DD), (**C**) initiation-to-initiation (II), and (**D**) initiation-to-division (ID) intervals. Left and middle panels show representative data from slow (MOPS + alanine, 0.25 h^−1^) and fast (RDM + glucose, 1.60 h^−1^) growth, respectively. Right panels show the adder slope (*k*) versus growth rate (⟨*λ*⟩). A slope of zero indicates perfect adder behavior. Data are presented as single-cell points (gray), binned averages ± SEM (red), and a linear fit (black line; slope *k*). Error bars (on right panels) represent 95% confidence intervals of the fitted slopes. *n* ≥ 1000 cells per condition. See appendix file 1 for details.

We quantified the adder behavior for three key intervals: division-to-division, initiation-to-initiation, and initiation-to-division (Fig. 1A). The likelihood of each adder correlation was measured by the slope (*k*) of a linear regression between the size at the start of the interval and the size added during it; a theoretical slope of zero indicates perfect adder behavior (Fig. 1B-D, fig. S2). Note that cell length was used as a proxy for cell size as previous literatures (*6, 17, 19*) (see Supplementary Materials). Our results revealed a clear hierarchy. The DD and II adders showed significant growth-rate dependence. The DD adder slope (*k*_*dd*_) decreased from 0.07 ± 0.08 at fast growth (⟨*λ*⟩ = 1.60 h^−1^) to -0.34 ± 0.06 under the slowest condition (⟨*λ*⟩ = 0.25 h^−1^), indicating a pronounced deviation from adder behavior (Fig. 1B). Conversely, the II adder slope (*k*_*ii*_) was near zero at slow growth (⟨*λ*⟩ < 1.0 h^−1^) but fell to -0.48 ± 0.04 at fast growth (⟨*λ*⟩ = 1.60 h^−1^, Fig. 1C). In contrast, the ID adder slope (*k*_*id*_) remained consistently close to zero across all conditions (Fig. 1D). This robust invariance of the ID adder was confirmed to be independent of the methods employed to quantify the slope (fig. S3A) and was further supported by concordance with re-analyzed data from previous studies (fig. S3B). Thus, systematic quantification demonstrates that only the ID adder provides a universal, growth-rate-invariant mechanism, directly coupling replication initiation to division.

### Inter-cycle correlations underlie growth-rate-dependent adder behaviors

The divergent growth-rate responses of the three adders suggest that the DD and II correlations may be emergent properties, rather than independent control mechanisms. We therefore investigated if statistical dependencies between successive added sizes, which we collectively term inter-cycle correlations (ICCs), could explain their behavior.

We first examined the ICC between initiation events (the inter-initiation correlation). The II adder itself is a predicted outcome of the well-established initiator-titration model, wherein DnaA accumulates to a chromosomal threshold to trigger replication (*28, 29*). This mechanism inherently yields near-perfect II adder behavior under non-overlapping replication cycles (slow-growth regime). However, at fast growth rates where replication cycles overlap, a key modulation occurs: chromosomes from two concurrent initiation events compete for the limited pool of free DnaA. This competition introduces a negative correlation between successive II added sizes (Δ_*ii*_).

Consistent with this picture, our data show this inter-initiation correlation (ICC of Δ_*ii*_) decreases from ∼0 to -0.23 as growth rate increases (Fig. 2A). This trend is quantitatively captured by various simulations of the titration mechanism under overlapping cycles (Fig. 2A, right; Supplementary Materials), which recapitulate the observed decline in *k*_*ii*_ (Fig. 2B). Thus, the growth-rate dependence of the II adder slope arises because the core titration mechanism is increasingly modulated by inter-initiation competition as replication overlap intensifies.

**Figure 2.**
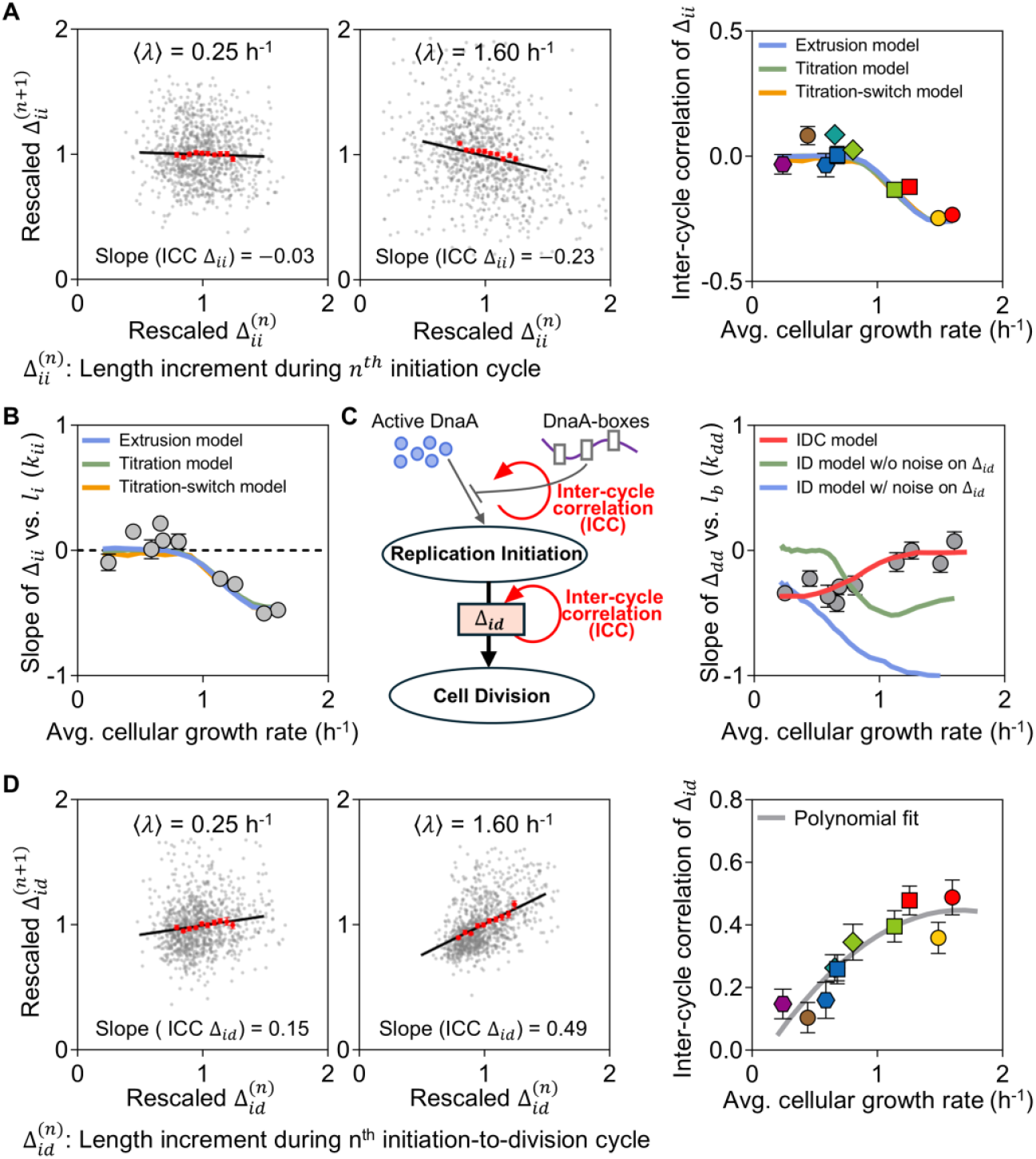
Growth-rate-dependent DD and II adder behaviors emerge from inter-cycle correlations. (**A**) Inter-cycle correlation between successive initiation-added sizes (ICC of Δ_*ii*_) for slow (left) and fast (middle) growth. Right panel shows ICC of Δ_*ii*_ versus growth rate, with predictions from various titration models (Supplementary Materials). (**B**) Numerical simulations of titration models recapitulating the growth-rate dependence of the II adder slope (*k*_*ii*_). (**C**) Illustration and numerical simulation of the IDC model (incorporating both ICC of Δ_*ii*_ and ICC of Δ_*id*_). The IDC model accurately predicts the DD adder slope (*k*_*dd*_) trend, unlike the ID-only model with or without Δ_*id*_ noise. (**D**) Inter-cycle correlation between successive ID-added sizes (ICC of Δ_*id*_) for slow (left) and fast (middle) growth. Right panel shows ICC of Δ_*id*_ versus growth rate. The empirical polynomial fit (gray line) was used as input for the IDC model in (**C**). In (**A**) and (**D**), data are presented as in Fig. 1. *n* ≥ 1000 cells per condition.

However, a model combining only this initiation-titration mechanism with the ID adder principle (the ID model, Supplementary Materials) fails, as it predicts a DD adder trend that contradicts our data (Fig. 2C, right, green and blue lines). This indicates that an additional source of correlation is missing. We reasoned that the initiation-to-division (ID) intervals themselves might be correlated. We measured the inter-cycle correlation between successive ID added sizes (ICC of Δ_*id*_) and found a positive correlation that increases with growth rate (Fig. 2D), contrasting sharply with the negative inter-initiation correlation (Fig. 2A). The biological origin of this positive inter-interval correlation is not yet known. To test its explanatory power, we incorporated this empirically measured correlation into a unified model (Initiation-to-Division Correlation, IDC model; Fig. 2C, left; Supplementary Materials). Strikingly, the model now accurately predicts the measured growth-rate dependence of the DD adder (Fig. 2C, right, red line). Thus, the seemingly complex and divergent behaviors of the DD and II adders naturally emerge from the robust, invariant ID adder and initiation titration mechanism combined with two key inter-cycle correlations. This suggests a clear causal hierarchy: the initiation titration and ID adder are the fundamental rules, whereas the DD and II adder correlations are derivative phenomena shaped by ICCs.

This framework fundamentally distinguishes our IDC model from the prior “double-adder” model (*19*), which posits two independent adders (II and ID). While both the double-adder and our IDC model posit the causal mechanism of ID adder, our model adopts the molecular level initiation titration mechanism and the experimentally observed ICCs, yielding a quantitative explanation of both (i) the growth-rate-dependent failure of the II adder and (ii) the full variation in the DD adder slope, neither of which the double-adder model adequately captures.

### Division mass is causally set by initiation mass

The robust single-cell correlation of the ID adder suggests a linkage between replication initiation and division. To test if this reflects a direct causal mechanism, we asked whether the same principle operates at the population level (Fig. 3A): if the population-averaged initiation mass (*M*_*i*_) is perturbed, does the population-averaged division mass (*M*_*d*_) change as predicted by the ID adder (Δ_*ID*_= *M*_*d*_ − *M*_*i*_ = constant)?

**Figure 3.**
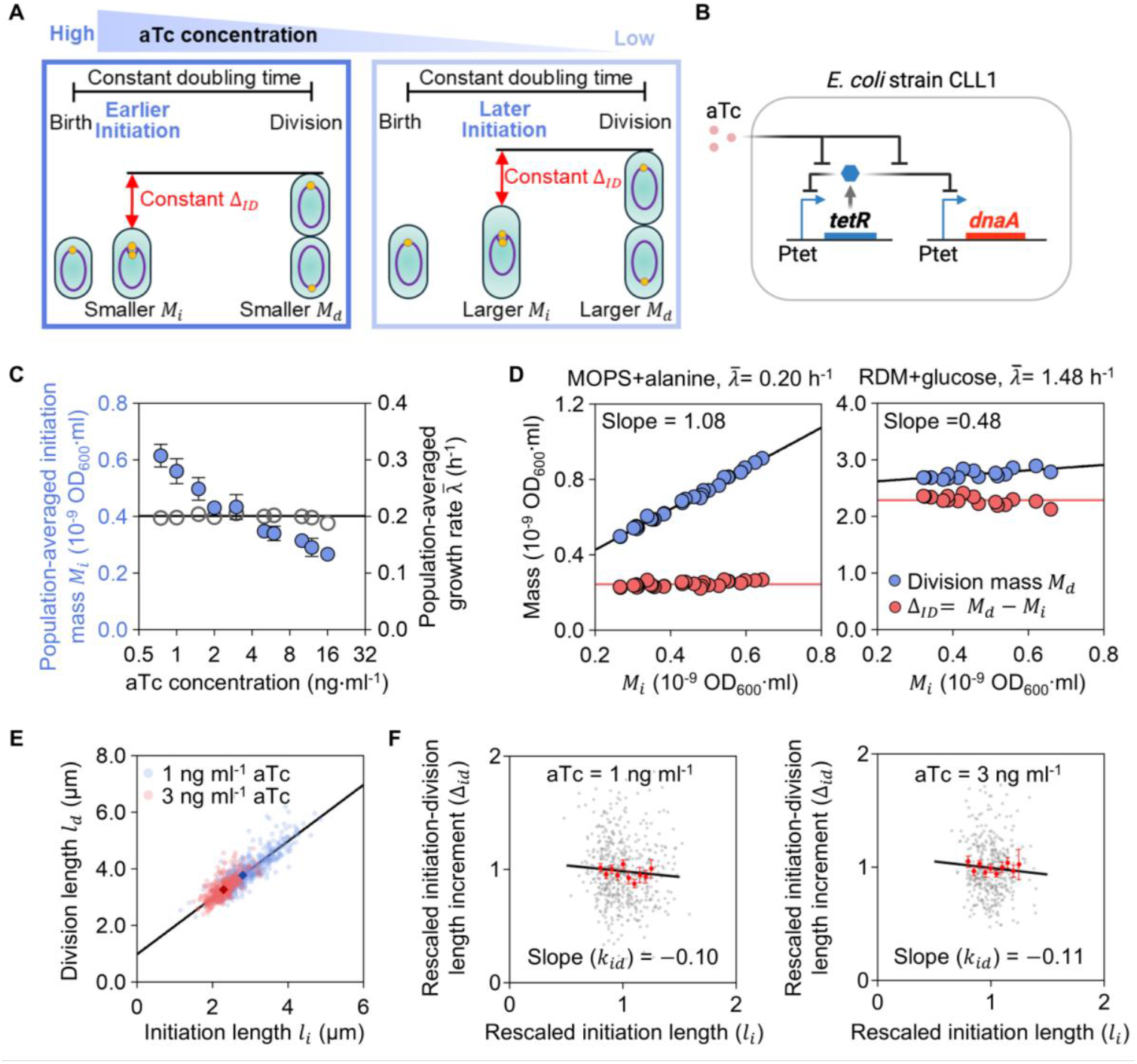
Perturbing initiation mass causally sets division mass via the ID adder. **(A)** Schematic of the experimental design. Modulating initiation mass (*M*_*i*_) via aTc is predicted to alter division mass (*M*_*d*_) while keeping the added mass (Δ_*ID*_) constant, according to the ID adder principle. **(B)** Genetic circuit of the DnaA-titratable strain (CLL1). **(C)** Population-averaged initiation mass (*M*_*i*_) and growth rate 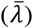 as functions of aTc concentration. **(D)** Relationship between *M*_*d*_ and *M*_*i*_ under slow growth (MOPS + alanine, 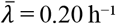, left) and fast growth (RDM + glucose, 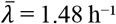, right). Blue dots, *M*_*d*_; red dots, Δ_*ID*_. Black lines, linear fits (slope indicated). Red lines, mean Δ_*ID*_ . Data in **(C, D)** are from ≥3 independent experiments. **(E)** Single-cell correlation between division length (*l*_*d*_) and initiation length (*l*_*i*_) for CLL1 with DnaN-YPet cultured in MOPS + alanine with 1 ng ml^−1^ (light red) or 3 ng ml^−1^ (light blue) aTc. Dark red/blue diamonds, population means. Black line, linear fit (slope = 1.00). *n* = 505 (1 ng ml^−1^); *n* = 386 (3 ng ml^−1^). **(F)** Single-cell ID added length (Δ_*id*_) versus *l*_*i*_ under the same conditions as in **(E)**. Data presentation as in Figure 1. *n* = 496 (1 ng ml^−1^); *n* = 377 (3 ng ml^−1^).

We specifically tuned *M*_*i*_ over a ≥2-fold range using a DnaA-titration strain (CLL1 in Table S1), wherein the expression of the replication initiator DnaA is controlled by a titratable promoter (Fig. 3BC, fig. S4A). Critically, by carefully selecting inducer concentrations, we achieved this perturbation without altering the growth rate (Fig. 3C, fig. S4B), ensuring that observed changes in division were not indirectly caused by global variations in metabolic flux.

Consistent with a departure from a simple timer, the average *C* + *D* period decreased as *M*_*i*_ increased (fig. S4C). More importantly, under slow-growth conditions 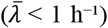, *M*_*d*_ increased linearly with *M*_*i*_ with a slope close to 1 (Fig. 3D, left; fig. S4D), whereas Δ_*ID*_ remained independent of *M*_*i*_ (CV < 6%) (Fig. 3D, left; fig. S4E). In the fast-growth condition 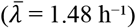, the same principle held, but its manifestation was shaped by its average cell mass (Fig. 3D; fig. S5). Here, the Δ_*ID*_ (∼2.29×10^-9^ OD_600_·ml) is substantially larger than *M*_*i*_ itself. Therefore, a similar ≥2-fold change in *M*_*i*_ resulted in a more modest relative change in *M*_*d*_ (Fig. 3D, right; fig. S5). The invariance of Δ_*ID*_ (CV < 5%) across this regime suggests that the underlying rule is the same (Fig. 3D, right). This one-to-one correspondence proves that initiation mass directly and causally sets the division size.

We next examined this relationship at the single-cell level. Cells were first grown to steady state at a low aTc concentration (1 ng ml^−1^), then shifted to a high concentration (3 ng ml^−1^) and allowed to reach a new steady state (fig. S6A–C). Under both conditions, division length scaled linearly with initiation length with a slope near unity (Fig. 3E), matching the population-averaged trend (Fig. 3D). Single-cell data tightly followed the population-level line (colored dots in Fig. 3E), and the single-cell ID adder slope (*k*_*id*_) remained close to zero across conditions (Fig. 3F; fig. S6D). These results indicate that the ID adder principle governs both single-cell behavior and population-level size control.

### Initiation-to-division hierarchy is preserved upon orthogonal reprogramming of division mass

To further test the causal influence of initiation mass on division mass at the population level, we asked whether this relationship holds when the absolute division mass is genetically reprogrammed. If initiation mass (*M*_*i*_) directly sets division mass (*M*_*d*_) as per the ID adder principle, then altering the expression of division proteins should systematically reset the division mass setpoint, while the added mass (*Δ*_*ID*_), and hence the causal link from *M*_*i*_ to *M*_*d*_, should remain intact.

We constructed a strain (CLL2 in Table S1) where the essential *dcw* gene cluster (including *ftsZ, ftsL*, and *murE*) is under the control of a titratable *P*_lac_-*lacI* circuit (Fig. 4A). Titrating with IPTG allowed us to fine-tune the population-averaged division mass (*M*_*d*_) over a >1.6-fold range (from ∼0.54 to ∼0.89 ×10^-9^ OD_600_·ml), while the population-averaged initiation mass (*M*_*i*_) and growth rate 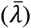 were unaffected (Fig. 4B, fig. S7A).

**Figure 4.**
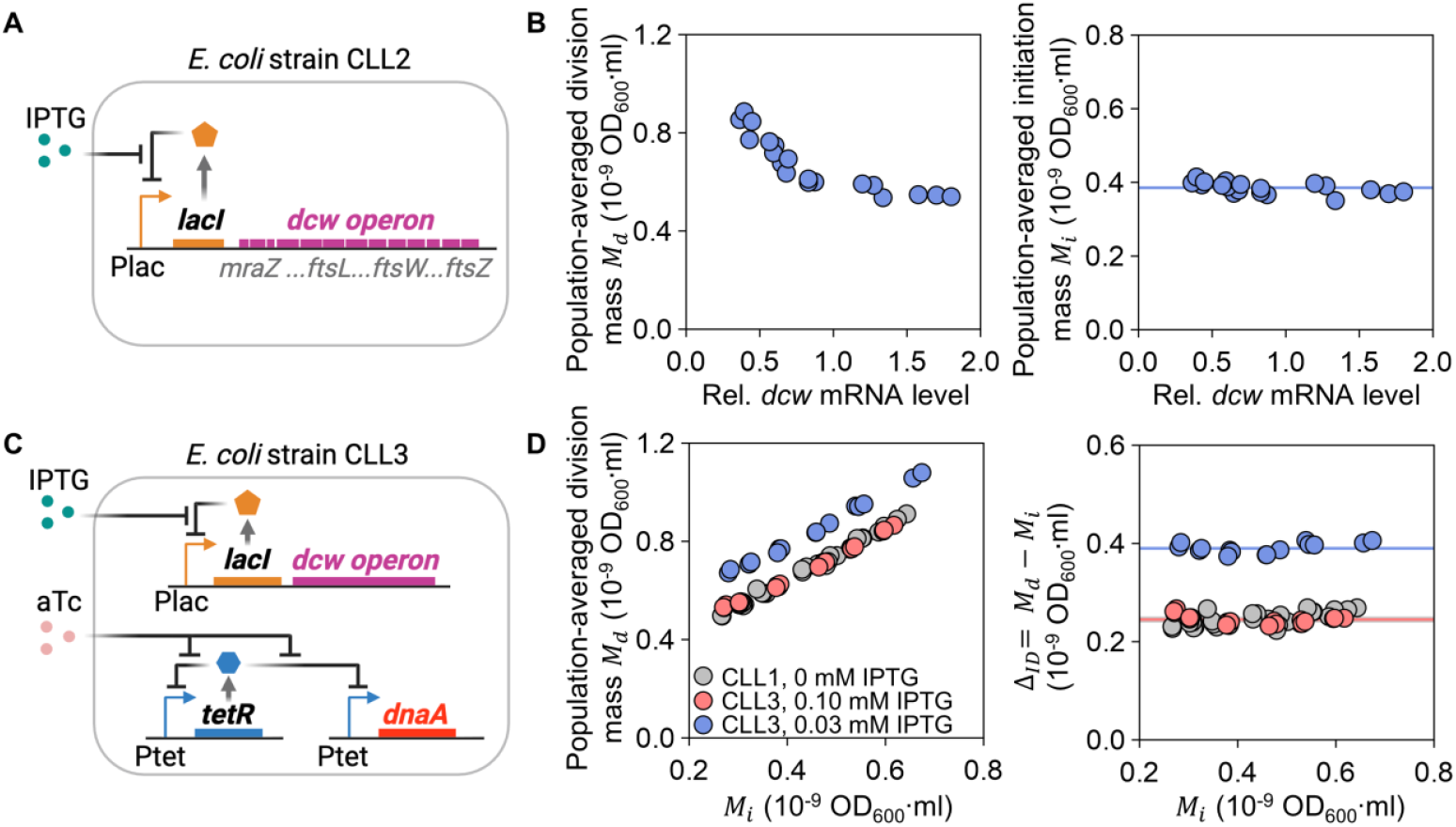
The ID adder principle is robust to genetic reprogramming of the division machinery expression. (**A**) Schematic of the *dcw*-titratable strain (CLL2), where division gene expression is controlled by IPTG concentrations. (**B**) Titrating *dcw* expression tunes division mass (*M*_*d*_, left) without affecting initiation mass (*M*_*i*_, right) in slow growth (MOPS + alanine). Blue line, mean. (**C**) Schematic of the dual-control strain (CLL3) for orthogonal perturbation of division and initiation. (**D**) Left: At fixed *dcw* expression, varying *M*_*i*_ in strain CLL3 proportionally shifts *M*_*d*_ while Δ_*ID*_ remains constant. Right: Reprogramming the division setpoint (by changing *dcw* expression level) resets the absolute value of Δ_*ID*_, but at each setpoint, Δ_*ID*_ remains invariant with *M*_*i*_. Colored lines indicate mean Δ_*ID*_ for each condition. Data are from ≥3 independent experiments.

To rigorously test the hierarchy, we engineered a dual-control strain (CLL3 in Table S1, Fig. 4C) by integrating the *dcw* and *dnaA* titration systems, enabling orthogonal control over division mass (*M*_*d*_) and initiation mass (*M*_*i*_). This approach yielded two pivotal findings. First, at any fixed division gene expression level, varying *M*_*i*_ proportionally shifted *M*_*d*_ (slope ≈ 1) whereas the ID added mass (*Δ*_*ID*_) remained constant (CV < 6%; Fig. 4D, left; fig. S7B). Second, and conversely, reprogramming *M*_*d*_ over a >1.2-fold range systematically reset the absolute value of *Δ*_*ID*_, yet at each new setpoint, *Δ*_*ID*_ remained invariant with *M*_*i*_ (Fig. 4D, right; fig. S7C). Thus, while the abundance of division proteins modulates the “how much” of mass addition, the initiation event governs the “when” according to the mass-additive rule.

### Cell division is independent of DNA replication termination

The traditional view that replication termination gates division (*1, 9, 21, 30-32*) is challenged here. We developed a stringent experimental system to isolate the effect of replication-termination timing. By constructing a *thyA deoB* double knockout strain (*33, 34*), we restricted the synthesis of dTTP, the DNA replication substrate, to exogenous thymine (Fig. 5A). Titrating thymine from 7 to 100 μg ml^−1^ allowed us to modulate the replication elongation rate in MOPS+glycerol, thereby varying the *C* period, the time from initiation to termination (from 1.08 to 1.28 hours) without altering the initiation mass or the bulk growth rate (CV < 5%) (Fig. 5B-D).

**Figure 5.**
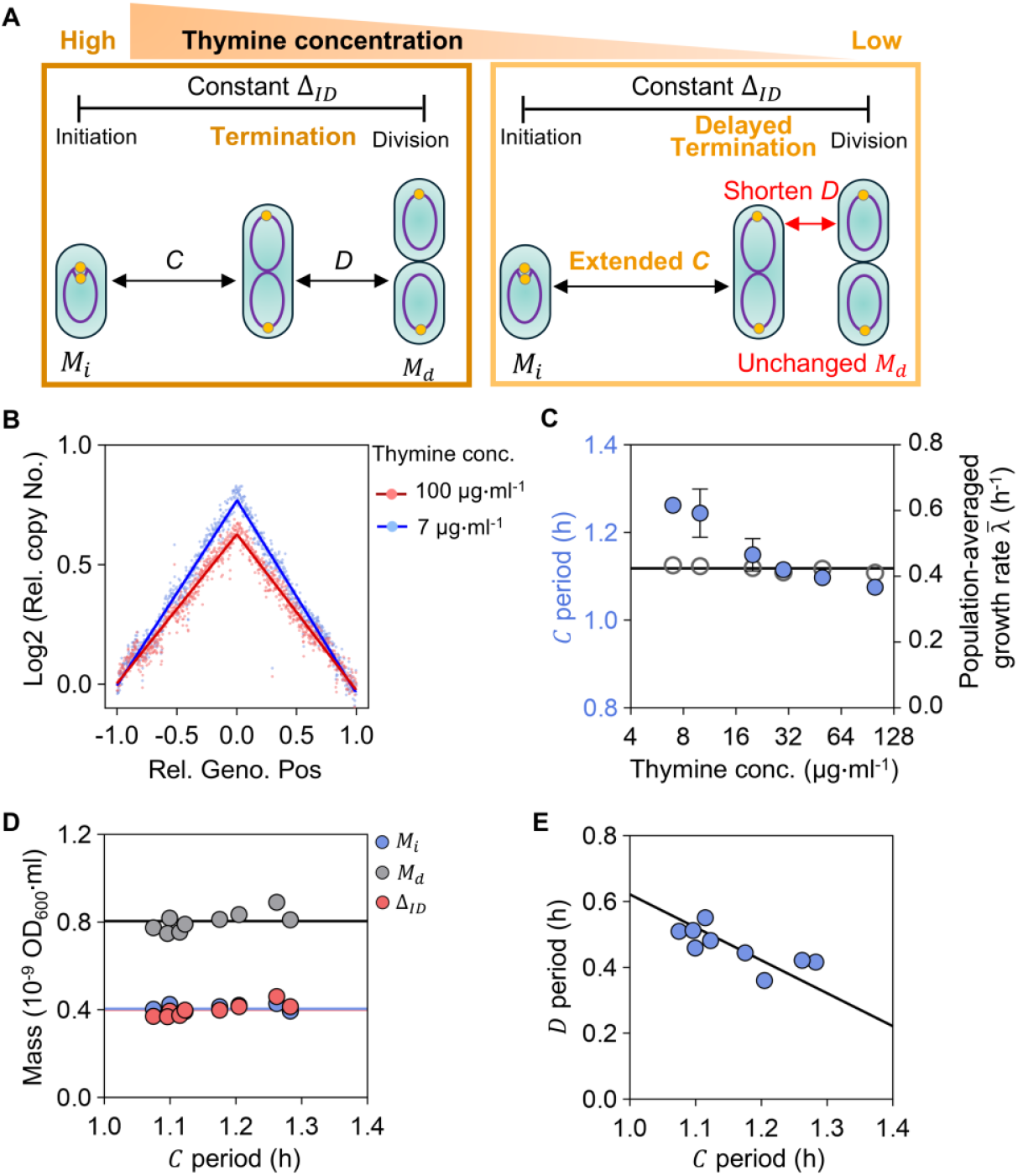
Cell division is uncoupled from replication termination. (**A**) Experimental principle: How division is affected by manipulating thymine concentration supplied to a thymine-requiring mutant thus altering *C* period without affecting bulk growth or initiation. (**B**) Chromosomal copy number profiles used to determine replication time *C* under high (100 μg ml^-1^) and low (7 μg ml^-1^) thymine concentrations. (**C**) The *C* period (blue) and mean growth rate 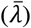 (gray) as functions of thymine concentration. (**D**) Initiation mass (*M*_*i*_), division mass (*M*_*d*_), and added mass (Δ_*ID*_) remain invariant across a range of *C* periods. Colored lines, mean values. (**E**) The *D* period shortens in a compensatory manner as the *C* period lengthens, preserving the ID-added mass. Solid line shows the fit ⟨*D*⟩ = ⟨ *C* + *D*⟩ − ⟨*C*⟩. Data in (**C**-**E**) are from ≥ 2 independent experiments.

We found that cell division is uncoupled from DNA replication termination. Across variations in the *C* period, the initiation mass (*M*_*i*_), the division mass (*M*_*d*_) and the ID added mass (Δ_*ID*_) remained invariant (CV < 8%; Fig. 5D). This invariance was maintained by a dynamic compensatory mechanism: the *D* period shortened as the *C* period lengthened, precisely preserving the constant Δ_*ID*_ dictated by the ID adder principle (Fig. 5E). Since division proceeds on schedule regardless of when termination occurs, we conclude that replication termination does not function as a checkpoint for the mechanisms underlying the ID adder principle. Instead, division timing is set at initiation, with the *C* and *D* periods serving as flexible intervals to fulfill the mass-additive rule (Fig. 5DE). Since division proceeds on time regardless of when termination occurs, we conclude that termination is not a necessary checkpoint for the ID-adder principle.

## DISCUSSION

For decades, understanding of the bacterial cell cycle has been divided between the classic timer model and various, often inconsistent, adder phenomena (*1, 19, 24, 35-37*). This study resolves these long-standing contradictions by establishing the ID adder as the central, causal, and hierarchically dominant principle. We demonstrate that cell division is directly coupled to DNA replication initiation through a mass-additive mechanism. Through causal perturbations, we show that the ID adder functions as a fundamental regulator, operating upstream of the expression level of the division machinery, which can modulate the absolute size setpoint without altering the underlying adder logic. This hierarchy ensures that the critical sequence of genome duplication followed by division is maintained, prioritizing genomic fidelity over mass precision, which is a clear evolutionary advantage.

The power of this principle lies in its ability to unify disparate observations. Our IDC model quantitatively shows that the growth-rate dependencies of the DD and II adders, long considered hallmarks of independent control, naturally arise from the initiation titration mechanism and the ID adder rule, coupled with a positive inter-cycle correlation between overlapping ID intervals. Consequently, these adder behaviors are not independent controls but emergent properties of a clear causal hierarchy: the ID adder constitutes the fundamental regulatory rule, while the II and DD adder correlations are phenomenological outcomes shaped by inter-cycle statistics under different physiological contexts (Fig. 2). This framework reconciles the apparent conflicts among the classic timer, single adders, and the double-adder model, resolving the paradox noted previously (*24*) wherein models like the double-adder could not coherently explain the full spectrum of adder behaviors.

Importantly, we do not claim the ID adder is an inflexible, solitary rule. Its dominance is bounded by physiological limits, defining a specific “adder regime” for cell cycle control. In wild-type cells, Δ_*ID*_ remains constant within a defined initiation mass range; beyond this threshold, whether by extreme initiation perturbations, severe division gene depletion, or thymine limitation, a regime shift occurs (fig. S8). Under such stress, division may become constrained by replication completion (fig. S8A), shift toward sizer-like behavior (fig. S8B), or exhibit increased added mass (fig. S8C), consistent with earlier studies (*21, 33, 38*). These observations delineate the operational boundaries of the ID adder and reveal a hierarchy of control strategies that ensure division fidelity under stress.

Notably, the altered division timing under extreme perturbations can be accounted for by incorporating concurrent cell cycles into our IDC model (Supplementary Materials), aligning with the broader conceptual framework of “concurrent cell-cycle processes” (*23, 39*), where multiple signals integrate to ensure robustness. Within this framework, our contribution is twofold: (1) we identify the ID adder as the replication-centric process that replaces the classical constant-time assumption, and (2) we establish it as the dominant principle under standard physiological conditions. The ID adder thus provides the primary mass-setting signal, onto which other checkpoints and stress responses are layered.

Seemingly conflicting past evidence that still awaits clarification was obtained regarding changes in the *D* period under thymine limitation in thymine auxotrophic strains. Using elution experiments (“baby machine”), longer *C* (at lower thymine concentration) results in shorter *D* (*40*), whereas multi-constricted spheroidal cells display a parallel rise in *D* (*41*), likely due to meager supply of FtsZ to complete ongoing division processes. Whether this discrepancy can be resolved by the framework established here remains an intriguing question for future investigation.

The molecular implementation of this robust principle presents a fascinating frontier. We propose that the starting and ending points of the ID interval, while observed as initiation and division, may encompass a broader physiological process. The starting point could be linked to the culmination of initiation events, such as the end of the eclipse period or the duplication of key *oriC*-proximal regions (*22, 26, 42*). Similarly, the endpoint may relate to the activation of the divisome or the onset of constriction. Within this framework, two non-exclusive mechanistic paradigms could instantiate the adder: an accumulation model (*6, 43*), based on the synthesis of a division component starting after initiation and triggers division at a threshold, or a spatial-ratiotic model, wherein a landmark established at initiation is monitored until a critical spatial increment threshold is reached to trigger division. This principle of coupling division to a replication-associated event also offers a new perspective for identifying the primary mechanism ensuring faithful genetic inheritance, moving beyond the paradigm of separate *C* and *D* periods gated by termination.

Given the fundamental nature of coordinating genome replication with division, the logic of the ID adder may represent a universally conserved strategy. In eukaryotic cells, where mass homeostasis is intricately linked to G1/S progression (*44, 45*), the principle of using replication initiation to gate a subsequent mass increment to division could represent a conserved, overarching logic, albeit implemented within a vastly more complex regulatory network. Thus, our work not only redefines the quantitative principles of the bacterial cell cycle but also provides a conceptual framework for exploring the deep logic of cell cycle control across the tree of life.

## Methods and models summary

A detail of experiments and the technical aspects of models are reported in the Supplementary Materials.

## Supporting information

Detailed Materials and Methods, Model Descriptions, and Supplementary Figures and Tables

## Acknowledgments

We thank Terence Hwa and Nancy Kleckner for valuable discussions and comments on the manuscript.

## Funding

National Natural Science Foundation of China 32025022 (C.L.)

National Natural Science Foundation of China 32230062 (C.L.)

Strategic Priority Research Program of the Chinese Academy of Sciences XDB0480000 (C.L.)

National Key R&D Program of China 2018YFA0902701 (C.L.)

Joint NSFC-ISF Research Grant 32061143021 (C.L.)

ISF-NSPC Joint Research Program 3320/20 (A.Z. & C.L.)

US-IL Binational Science Foundation 2017004 (A.Z.)

National Natural Science Foundation of China 32170042 (H.Z.)

Youth Innovation Promotion Association CAS 2022369 (H.Z.)

National Natural Science Foundation of China 31900024 (Z.Z.)

National Natural Science Foundation of China 32571666 (Y.B.)

National Key R&D Program of China 2024YFA0919600 (X.F.)

Shenzhen Science and Technology Program ZDSYS20220606100606013 (X.F.)

National Natural Science Foundation of China 31971350 (S.H.)

## Author contributions

C.L. conceived and supervised the study. C.L., A.Z., and H.Z. jointly devised the experimental protocols. Y.B., H.Z., and X.F. collaborated on model development. D.L., H.Z., Q.C., W.H., and S.H. conducted the experiments. Z.Z. generated the analysis code and performed single-cell data analysis. Y.B. authored the simulation code and performed simulations. All authors collectively analyzed the results and contributed to manuscript writing.

## Data and materials availability

The calculated slopes of the single-cell correlations are provided in the Supplementary Materials. Simulation data can be generated using the custom-made code and the parameter sets provided. The code that was used for the simulations in this study are available at: https://github.com/BaiYangBqdq/initiation_to_division_adder. Further information and requests for resources and reagents used in this study can be requested from the lead contact Chenli Liu

