## Supplementary material for "DNA replication initiation causally sets the added mass for cell division in *Escherichia coli*": Detailed Materials and Methods, Model Descriptions, and Supplementary Figures and Tables

#### The PDF file includes:

1. Materials and Methods
2. Model descriptions
3. Figures S1 to S8
4. Table S1
5. References

#### Other Supplementary Material for this manuscript includes the following:

Figures S9 to S23
Tables S2 to S4

### 1. Materials and Methods

#### 1.1. Bacterial strains

All the strains used in this study were derived from the wild type *Escherichia coli* K12 strain AMB1655 (*1*). To visualize replication initiation, we fused the gene coding for fluorescent protein mScarlet-I (a gift from the Jin lab (*2*)) to the 3' end of the gene for the endogenous SeqA, employing a GGGS linker (*3*), accomplished through the CRISPR-Cas9 genome editing system (*4*). This procedure was performed in strain AMB1655, previously modified to lack *fliC*, thus creating strain ZH101.

To control the initiation mass of the cells, we constructed a *dnaA*-titratable strain (CLL1), by integrating the $P_{tet}$-*dnaA* construct proximal to the *oriC* locus, while a negative feedback loop, $P_{tet}$-*tetR*, was introduced independently at the *intS* locus. Subsequently, the native *dnaA* gene was substituted by a kanamycin-resistant gene. To monitor replication initiation events of the *dnaA*-titratable strain at the single-cell level, we introduced the DnaN-Ypet marker following a previously described method (*5*) and deleted the *fliC* gene (CLL1N). To perturb the expression level of the division and cell wall (*dcw*) gene cluster, we constructed a *dcw*-titratable strain (CLL2) by replacing the native $P_{mraZ}$-containing upstream regulatory region (−461 to −21 bp relative to the *mraZ* start codon) with an inducible $P_{lac}$–*lacI* cassette. The same edit was incorporated into CLL1, allowing orthogonal control of division mass and replication initiation respectively. In parallel, to modulate the replication time $C$ via thymine availability (*6*), *thyA* and *deoB* were deleted in the *E. coli* K12 strain AMB1655.

Information for all strains are listed in Table S1

#### 1.2. Growth conditions

For stain construction, the cells were cultured in Luria-Bertani medium (LB) at 37 °C. Plasmids were maintained with appropriate antibiotics: kanamycin (50 $\mu g \cdot ml^{-1}$), spectinomycin (50 $\mu g \cdot ml^{-1}$), and chloramphenicol (25 $\mu g \cdot ml^{-1}$). For specific strains, we used the following inducers and supplements: the *dnaA*-titratable strain was induced with 10 $ng \cdot ml^{-1}$ anhydrotetracycline (aTc); the *dcw*-titratable strain was induced with 0.25 mM Isopropyl β-D-1-thiogalactopyranoside (IPTG), and the double knockout Δ*thyA* Δ*deoB* strain was sustained with 50 $\mu g \cdot ml^{-1}$ thymine.

In single-cell time-lapse experiments, we used MOPS media supplied with different carbon sources and amino acids, as listed in Figure 1 and detailed previously (*7*). Notably, for the *dnaA*-titratable strain, we cultured it in MOPS + alanine medium containing 1 $ng \cdot ml^{-1}$ aTc, loaded it into the mother machine chip, then monitored it for 24 h under 1 $ng \cdot ml^{-1}$ aTc to collect single-cell data under steady-state conditions, followed by switching to 3 $ng \cdot ml^{-1}$ aTc for another 36 h observation.

In initiation mass perturbation experiments, CLL1 cells were grown in RDM+glucose (0.4% w/v), MOPS+EZ+glycerol (0.4% v/v), MOPS+glucose (0.4% w/v), MOPS+glycerol (0.4% v/v) and MOPS+alanine (40 mM), each supplemented with varying concentrations of aTc. Experiments were performed at 37 °C with shaking at 150 rpm, and steady-state growth was monitored, maintained and verified as previously described (*7*), with refined dilution parameters as necessary. All Reagents or resources used in this study are listed in Table S2.

##### 1.3. Microfluidics

In all single-cell experiments, *E. coli* was cultured in a 'mother machine' microfluidic device for at least 20 generations to maintain steady-state growth, during which growth and replication were monitored. Three different trap depths, including 0.8 µm, 1.0 µm, and 1.2 µm, were employed to accommodate cells with varied widths in different growth media (*8*). Throughout the single-cell experiments, fresh medium was perfused through the device using a pressurized pumping system at a constant pressure of 400 mbar.

##### 1.4. Cell preparation for single-cell experiments

A single colony from an LB agar plate was inoculated in 2 ml of liquid culture, which was grown overnight at 37°C with shaking at 150 rpm. This pre-culture was diluted into fresh medium by 1:500 when the $OD_{600}$ exceeded 1.0. Cells were then serially reinoculated four more times in a 37°C water bath, each time at an $OD_{600}$ of ~0.2, to ensure steady-state growth prior to microfluidic loading. The growth rate of batch-cultured cells was monitored by measuring $OD_{600}$ using a spectrometer (Genesys 10s, Thermo Fisher Scientific).

Prior to loading, the microfluidic device was treated with surfactant Pluronic F127 (Merck) to prevent bacterial adhesion to the surface of main channels. Cells were harvested and concentrated by centrifugation, loaded into the device using a 2-ml syringe, and briefly spun at 2000 rpm to settle them into the side channels. The assembled mother machine and medium reservoir were kept at 37 °C for one hour with medium perfusion, before time-lapse imaging commenced, and this setup was maintained throughout the experiment.

##### 1.5. Microscopy and image acquisition

The detailed setup of the microscope was described in (*9*): Image acquisitions were performed on an inverted microscope (IX-83, Olympus) with 100x oil objective (Olympus), an automated XY-stage (ASI, MS2000) and a sCMOS camera (Prime BSI, photometrics). Fluorescence images were acquired with solid-state lasers (Coherent OBIS: 405–100 LX, 561–100 LS). Acquisition intervals varied across growth media but were set to ensure more than 12 frames per generation under all conditions.

##### 1.6. Cell segmentation and lineage reconstruction

Phase-contrast images were segmented using a custom MATLAB program. First, the side channels of the mother machine device were registered and extracted. Cells within these channels were then identified using a deep-learning model adapted from DeLTA (*10*). Cell meshes were generated from the resulting masks by applying algorithms adapted from Oufti (*11*).

Cell length was determined from the skeleton of these meshes. The division cycle was defined as the period from a cell's birth until immediately after its subsequent division. The division length was calculated as the sum of the lengths of the two newborn daughter cells. Cell lineages were reconstructed by tracking cell position and length between consecutive frames, and growth rate for each cell was obtained through exponential fitting.

##### 1.7. Replisome foci analysis

To visualize DNA replication, the replisome was fluorescently tagged by fusing SeqA with mScarlet-I. Fluorescence images were first aligned to their corresponding phase contrast images. Emergence, disappearance, and location of seqA-mScarlet spots were used to evaluate replication initiation in all growth rates. Replisome foci were detected using an algorithm adapted for low signal-to-noise ratio images, which employs the LL-ratio to determine the location and number of foci per cell (*12*).

In conditions without overlapping replication cycles, new initiations were identified by the emergence of new foci adjacent to their ancestor generation. During overlapping replication, a single pair of replisomes may undergo several cycles of splitting and rejoining before a new, permanent initiation event occurs. This pattern allowed us to construct initiation-to-initiation lineages and test the II-adder phenomenon across growth conditions. Specifically, we designed GUI programs to inspect the results from both channels and lineages, in order to fix or remove possible mistakes.

Before cell division, the sister chromosomes from a specific replication round segregate into the daughter cells. Linking each replication-initiation to its corresponding division allowed us to replication-division lineages and calculated the added length between them (ID-adder). As cells may divide once or more times within this interval, division mass was calculated precisely by accounting for the length ratio between daughters for each division, rather than assuming symmetrical division.

##### 1.8. Calculation of the slopes

To test the adder principle in various growth conditions, we analysed the relationship between the added mass (Y-axis) and the initial mass (X-axis) for each process. Both parameters were first rescaled by their respective mean values. Outliers were defined as data points beyond three times standard deviations from the mean. The data were then binned along the X-axis, and bins containing fewer than 0.5% of the total samples were discarded. The analysis was performed using the center position of each remaining bin on the X-axis and the average or median Y-value. The same rules were applied in the calculation of inter-generation correlations.

The slope of this relationship was calculated using six distinct methods:

1. Linear regression on all data, excluding outliers.
2. Linear regression on all data.
3. Linear regression on the binned data (average or median), excluding outliers.
4. Linear regression on the binned data (average or median).
5. The covariance of all data, excluding outliers.
6. The covariance of all data.

Here, each calculation of slope was based on more than 1000 sample points. To illustrate the correlations in figures, we randomly chose 1000 sample points (gray scatters), plotted the mean and SEM of each bin (red dots), and the linear regression of all sample points without outliers (black line).

#### 1.9. Measurements of population-averaged parameters

Population-averaged growth rate ($\bar{\lambda}$), cell mass ($\bar{m}$) and cellular *oriC* number ($\bar{o}$) were characterized as previously described (*7*). Briefly, we pre-cultured cells for approximately 10 generations in exponential growth. We then carefully measured the $OD_{600}$ of the culture at assigned consecutive time points using a spectrometer (Genesys 10s, Thermo Fisher Scientific). The growth rate, i.e., inverse of the doubling time $\tau$, was calculated by fitting the data to an exponential curve. When the $OD_{600}$ reached approximately 0.15, samples were taken to quantify the $\bar{m}$, and $\bar{o}$; $\bar{m}$ was characterized by dividing the $OD_{600}$ by cell number concentration, which was measured using a flow cytometry (CytoFLEX, Beckman Coulter Life Sciences). Classical run-out experiment (*13, 14*) was used to characterize $\bar{o}$ as previously described (*7*), with the exceptions that no Triton X-100 have been used in the stain buffer and the final concentration of rifamycin was 500 $\mu g \cdot ml^{-1}$ for this study.

To quantify the expression levels of *dnaA* and the *dcw* gene cluster, we performed quantitative real-time RT-PCR (qPCR) as described previously (*15*). Briefly, total RNA was isolated from steady-state cultures (genomic DNA removed), reverse-transcribed into cDNA, and quantified with SYBR Green chemistry. Gene expression levels were normalized to the reference gene *rpoA*. Transcript levels of *mraY* (representing the *dcw* cluster) and of *dnaA* were measured and used as a proxy for cluster-wide expression, while *dnaA* transcript levels were measured in parallel. Relative expression levels of *dnaA* and *dcw* were calculated by normalizing to the expression level measured in the wild-type strain grown in the same medium. Primer sequences were as follows: *dnaA* (Forward: TACCCAATCGAGGACAAAAC; Reverse: CCCGATTGCAGGATGA GTT), *mraY* (Forward: CCGGGGAACTGGTTATTGTCT; Reverse: TTTCTA CCACGAACACGCCC), and *rpoA* (Forward: CTTCTTTGGTGCTGTACTCA; Reverse: TGGTTGATATCGAGCAAGTG)

The $C$ period was determined using a genome-wide marker-frequency approach based on deep sequencing as previously described *(3)*. Under steady-state growth, the population-averaged copy number of a chromosomal locus depends on its normalized genomic position (*oriC* = 0, *terC* = ±1), the $C$ and $D$ periods, and the mass doubling time (τ). Genomic DNA was extracted from steady-state cultures and subjected to high-depth sequencing. The chromosome was partitioned into evenly spaced genomic bins, and the read coverage of each bin was normalized to the terminus region (*terC*). Run-out samples, which exhibit uniform copy number across all loci after completion of ongoing replication, were used as controls for normalization. For each condition, $\log_2$(relative copy number) was plotted against the normalized chromosomal coordinate (m), and linear regression was performed separately for each of the two replication arms. The absolute slopes were averaged and used together with the measured doubling time to compute the $C = |k| \times \tau$, where $k$ denotes the average slope of the linear fit.

#### 2. Model descriptions

##### 2.1. Construction of the stochastic titration model

To understand the growth-rate dependency of the DNA initiation-initiation (II) adder, we revisited the classic initiator titration model (*16, 17*). This model is built on the principle that DNA replication initiation in bacteria is governed by the accumulation of the free active form of the initiator protein, DnaA and its titration of it through DnaA binding sites (DnaA-boxes) on the chromosome. Due to autoregulatory feedback on the *dnaA* promoter, the concentration of DnaA is maintained at a relatively constant level during steady-state growth (*18, 19*), As a result, the total cellular amount of DnaA scales proportionally with cell volume, which increases exponentially over time. In contrast, DnaA-boxes and their copy number is directly tied to the chromosomal replication state. Each initiation event doubles the number of *oriC* copies, and thus the number of DnaA-boxes, introducing demand for DnaA binding (*20, 21*). Following initiation, free DnaA gradually re-accumulates through growth-coupled synthesis until it reaches the threshold required to trigger the next round of initiation. This dynamic interplay between exponentially accumulating DnaA and stepwise-increasing DnaA-box numbers forms the basis for the robust replication initiation.

###### 2.1.1. Model construction

The classic titration model provides dynamical regulations of DNA replication initiation, but to simulate the single cell correlations between successive initiations, it is necessary to implement the stochastic single-cell model to simulate bacterial cell cycle and to parameterize properly across growth rates.

Central to the model is the titration of free DnaA proteins through DnaA-boxes on chromosomes. For simplicity, the DnaA boxes were evenly distributed on the chromosome while the chromosome structure is dynamically changing during cell cycle as DNA replicates (*16*). Such dynamics of the chromosome structure and thus number of DnaA boxes brings complexity to the continuous calculation so that we simulate it numerically. To numerically simulate the chromosome structure, a vector of DNA gene copies was created so that its length represents a linearly distributed chromosome while each element in this vector represents the copy number of the genes located on the chromosome at each site. It is then easy to calculate the total DNA and the number of DNA boxes according to the gene copies. The replication of the DNA is set by doubling the gene copies at proper locations at each time step.

The simulation tracks this process over time with a short step, updating cell volume, DnaA concentration [DnaA], and DNA copy number vector. Upon initiation, replication forks progress along the chromosome and terminate after the $C$ period. The stochastic was introduced into the gene expression level of DnaA at each initiation cycle with a Gaussian distributed noise. For each growth condition, the simulation is repeated multiple times to generate statistical ensembles of initiation mass and time.

##### 2.1.2. Parameterization

The model explores a range of growth rates, computing for each the corresponding $\tau$ and $C$, using empirically derived relationships. It simulates the bacterial cell cycle by integrating key physiological and molecular parameters that govern cell growth, DNA replication initiation, and division timing. The primary control variable is the growth rate ($\lambda$), which spans a biologically relevant range (0.1 to 2 $h^{-1}$), resulting of different nutrient conditions. The DNA replication duration $C$ is derived as a function of the growth rate using an empirical formula: $C = \left(\frac{\ln 2}{\lambda}\right) \times (0.8801 \times \lambda \times 60 + 0.2375)$ (*7*).

A central regulatory mechanism in the model is the accumulation of DnaA, which triggers the initiation of DNA replication upon exceeding a threshold. The DnaA synthesis rate is controlled by the parameter $\alpha$ that is set proportional to the growth rate and is fitted by the average initiation mass experimentally measured over growth rates (*7*) (Fig. S9, Table S3). The number of DnaA-boxes, denoted Abox, depends on inhibitory elements: the number of DnaA-boxes on *datA* site (denoted $nbox_{datA}$) and the number DnaA-boxes linearly distributed on the chromosome (denoted $nbox_{chrom}$). While $nbox_{datA} = 250$ and $nbox_{chrom} = 150$ in the same order of experimental reports (*17, 20, 21*). To account for biological variability, the model introduces stochasticity through the noise parameters: $\epsilon_A$ adds noise to DnaA synthesis rate. The noise level $\epsilon_A = 0.15$ to coordinate the experimental noise level on initiation mass.

For efficiency, 200 independent cells are simulated over 100 doubling times (DT) for each growth rate condition ($t_{max} = 100$ DT) with a fine time resolution ($dt = 0.2$ min) to accurately capture fast molecular dynamics. The outputs, including initiation, termination, and division events, are summed and used to compute correlation slopes and infer the dominant mass regulation mechanism.

In summary, the model uses a combination of deterministic scaling laws (e.g., growth rate dependence of $\tau$ and $C$ period) and stochastic elements (noise in DnaA, growth, and mass increments) to simulate realistic single-cell behavior. Parameters are carefully chosen to reflect known biological constraints and to enable systematic investigation of how bacteria maintain mass homeostasis and coordinate DNA replication with cell division across diverse growth conditions.

##### 2.1.3. Model implementation

The model is implemented with a matlab code as the following pseudocode:

**Variables:**

| | |
|---|---|
| $V$ | : Cell volume |
| $A$ | : DnaA protein amount per cell |
| $Abox$ | : DnaA-boxes number per cell |
| $DNAcopy$ | : 1D array representing gene copies on chromosome |
| $DNA$ | : Total DNA amount per cell |
| $OriC$ | : Chromosome replication origins per cell |
| $N_{datA}$ | : Copy number of *datA* per cell |
| $nrep$ | : Number of replications round counter |
| $Prep$ | : Positions of active replication fork |

**Parameters**

λ : Cell mass growth rate
$DT = \ln(2)/\lambda$ : Doubling time, $\tau$
$TT = 100DT$ : Total simulation time
$dt = 0.2$ min : Time step
$C$ : Duration of DNA replication (computed from $\lambda$)
$\alpha$ : DnaA synthesis rate
$\alpha_0$ : Expected DnaA synthesis rate (fitted value)
$nbox_{datA} = 250$ : Number of DnaA binding sites at *datA*
$nbox_{chrom} = 150$ : Number of DnaA-boxes per chromosome
$\epsilon_A = 0.15$ : Noise level on DnaA expression rate
$[A_f^c] = 0$ : Threshold for DNA replication initiation
$AboxRange = 0.8$ : The range of DnaA-boxes linearly distributed on chromosome

**While $t < t_{max}$ do**

- Update cell state:

  # Cell mass growth and DnaA synthesis
  $V = V + \lambda \cdot V \cdot dt$
  $A = A + \alpha \cdot V \cdot dt$
  # Get gene amounts and number of DnaA-boxes
  $DNA = sum(DNAcopy(1\colon AboxRange))$
  $N_{datA} = DNAcopy(1)$
  $OriC = DNAcopy(1)$
  $Abox = N_{datA} \cdot nbox_{datA} + DNA(1\colon AboxRange) \cdot nbox_{chrom}$

- DNA replication initiation:

  **If** $(A - Abox)/V > [A_f^c]$

  # Start a new replication fork at origin of the DNAcopy vector and double it
  $nrep = nrep + 1$
  $DNAcopy(1) = 2 \cdot DNAcopy(1)$
  $Prep(nrep) = 1$
  # Reset stochastic DnaA synthesis rate:
  Draw a random unit Gaussian variable $\eta$,
  $\alpha = \alpha_0 + \epsilon_A \eta$
  # Record cell volume, initiation time and origin number at initiation
  $V_i(nrep) = V/OriC$
  $T_i(nrep) = t$
  $O_i(nrep) = OriC$

  **end**

- Update DNAcopy:

  For each active fork in Prep:

**If** fork $Prep(nrep) \leq DNA\,length$:
$Prep(nrep) = Prep(nrep) + 1$
$DNAcopy(Prep(nrep)) = 2 \cdot DNAcopy(Prep(nrep))$
**else**
### Replication complete
$Prep(nrep) = Null$
**end**
**end**
Output: $V_i, T_i, O_i$

#### 2.2. Results of the stochastic titration model

##### 2.2.1. The growth dependent II adder

Using given growth rates, we run the simulation and calculate the mass per *oriC* over time, which increases exponentially and suddenly drops by half at the time of DNA replication initiation where *oriC* number doubles (Fig. S10AB). From the recorded $V_i^n$ and $T_i^n$, we performed a correlation analysis on the simulated data to distinguish between three canonical mass control strategies: **sizer**, **adder**, and **timer**. This analysis focuses on the relationship between the initiation mass in one cycle and key variables in the subsequent cycles—specifically, the next initiation mass, the volume added between initiations, and the time interval between initiations. From these, we compute the inter-initiation volume increment ($\Delta V_i$) and time interval ($\tau_{ii}$). The control strategy is determined by plotting the rescaled initiation mass in cycle *n* (Rescaled $V_i^n$) against three quantities: the initiation mass in cycle *n+1* (Rescaled $V_i^{n+1}$ ), where a slope near zero suggests a **sizer**; the added volume (Rescaled $\Delta V_i$), where a slope near zero indicates an **adder**; and the time between initiations (Rescaled $\tau_{ii}$), where a slope near zero suggests a **timer** (Fig. S10CD). Linear fits to the rescaled data provide quantitative evidence for which principle dominates the initiation timing control.

Across a range of growth rates from 0.1 to 2 $h^{-1}$, the II sizer slope (Fig. S11A) remains consistently positive and close to 0.5. In contrast, the II adder slope (Fig. S11B) decreases from near zero at slow growth to approximately –0.5 at fast growth, suggesting a transition from an adder-like behavior at low growth rates to a more complex regime where the volume added between initiations becomes negatively correlated with initial mass. This implies that in fast growth condition, a cell initiating at larger mass may add less volume between initiations. Finally, the II timer slope (Fig. S11C) is consistently negative, approaching –1 at high growth rates, indicating that inter-initiation time becomes increasingly dependent of initiation mass and instead follows a fixed or nearly fixed interval, especially under fast growth conditions.

##### 2.2.2. Overlapping cycle induces additional correlation

In this model, the total amount of DnaA represents cellular biomass, whereas the number of DnaA-boxes reflects the amount of DNA. Biomass grows exponentially (the gray line in Fig. S12) while DNA amount (DnaA-box No., the blue line in Fig. S12) increases linearly with a speed defined by the number of ongoing replication fork(s) and the replication speed per replication fork. The cell initiates a new round of DNA replication (orange solid dots in Fig. S12) when the biomass catches up the DNA amount, i.e., the amount of DnaA proteins catches up the number of DnaA-boxes.

Under slow-growth conditions, where the replication cycle is not overlapped, the classic titration model predicts adder correlation between initiations (*22*). The DnaA-box number increases as DNA replication initiates and remains constant after replication terminates. Therefore, the DnaA-box number per chromosome sets a fixed threshold for the amount of added DnaA protein per *oriC* between successive replication initiations. Remarkably, this threshold, and the amount of the added mass, remains independent of the cellular mass at the time of the previous initiation, thus leading to the II adder phenomenon (Fig. S12A).

In contrast, fast-growth conditions introduce overlapping DNA replication cycles, wherein a new round of chromosome replication initiates before the previous one terminates, leading to dynamically shifting in the number of added DnaA-boxes between consecutive replication initiations. Specifically, if the previous initiation was headed, i.e. smaller initiation mass, the distance between successive replication folks on a chromosome gets smaller, then the co-existence period of them becomes longer. So that more genome, and also more DnaA-boxes, were synthesized during two replication initiations, which sets a larger threshold for the DnaA proteins, and also the cell mass to accumulate. For the inversed case, retarded previous initiation (larger initiation mass) will result in less co-existence period of multiple replication folks on one chromosome and then result in smaller threshold (less new synthesized DnaA-boxes) for the added cell mass for the next initiation (Fig. S12B).

To further confirm the overlapping effect on II adder correlations, we decoupled the DNA replication time $C$ and the cell mass doubling time $\tau$. Because the ratio of the average time duration for a round of chromosome replication ($C$) and the average time duration between successive chromosome replication initiations ($\tau$) determined the overlapping strength of DNA replication initiation. By changing the $C$ at given $\tau$ or inverse, we simulated the initiation cycle, and observed that the II adder slope $k_{ii}$ changes only with the ratio $C/\tau$ (namely, the number of replication positions (*8*), Fig. S13A).

At various growth conditions, the ratio $C/\tau$ changes naturally with growth rates (Fig. S13B). With a linear function experimentally measured, we get the growth rate dependent slope as a function of $C/\tau$. As predicted, this inter-generational correlation of $\Delta_{ii}$ was observed experimentally (Figure. 2B). These results indicated that the interplay between chromosome replication and biomass growth might be responsible for the disruption of II adder behaviors in fast-growth regime.

##### 2.2.3. Effects of DnaA synthesis and DnaA-boxes amount

The key parameters regulating initiation mass are the DnaA synthesis rate and the number of DnaA-boxes per chromosome. Simulations at two different growth rates (doubling times of 30 min and 60 min) show that while these parameters shift the average initiation mass, they have only minor effects on adder correlations. Perturbing either parameter does not significantly alter the adder behavior (Fig. S14).

##### 2.2.4. DnaA-boxes distribution changes II adder

Regulation of the II adder principle is highly sensitive to the chromosomal distribution of DnaA-boxes, particularly under fast growth conditions. This sensitivity depends on two key factors: the number of DnaA-boxes at the *datA* locus and the spatial extent over which DnaA-boxes are distributed along the chromosome—quantified by the dimensionless parameter, *AboxRange*, which ranges from 0 to 1 and defines the fraction of the chromosome spanned by these sites.

As shown in Fig. S15AB, under non-overlapping replication conditions (doubling time DT = 60 min), variations in either the number of *datA* DnaA-boxes or their chromosomal distribution range have negligible effects on the II adder slope. However, under overlapping replication cycles (DT = 30 min), these parameters significantly influence the adder behavior. Quantitative analysis reveals that broadening the DnaA-box distribution—either by reducing the number of boxes at *datA* or by spreading them over a larger chromosomal region—drives a deviation from ideal II adder behavior.

Across a broader spectrum of growth rates, the *datA* DnaA-box number and the distribution range modulate the relationship between the II adder slope and average cellular growth rates in distinct ways (Fig. S15CD). Specifically, decreasing the number of DnaA-boxes at *datA* amplifies the magnitude of deviation from the II adder behavior under fast growth, without shifting the critical growth rate at which the adder slope begins to deviate from zero.

#### 2.3. Other models for DNA replication initiation regulation

In this section, two major extensions of the classic titration model are described and implemented to stochastic version over growth rates.

##### 2.3.1. The titration-switch model

The titration-switch model extends the classic initiator titration framework by incorporating key biochemical and genomic features of DnaA regulation to more accurately capture the dynamics of DNA replication initiation. At its core, the model notes that DnaA exists in two forms: an active DnaA-ATP state capable of promoting initiation and a less active DnaA-ADP form. The active DnaA pool accumulates in proportion to cellular growth and is depleted through binding to high-affinity sites across the chromosome at regulatory loci like *datA* and RIDA (regulatory inactivation of DnaA). The DnaA-ATP was also reactivated by acidic lipid and DARS1/2 locus. Integrating the molecular titration and activity regulation within a dynamically replicating chromosome provides a mechanistic basis for robust, growth-rate-adapted control of replication-initiation timing.

To simulate this model, we followed the titration model to determine the total number of high-affinity DnaA binding sites (A-boxes) available for sequestration, and incorporated the switch effect between DnaA-ATP and ADP formulated by Wolde *et al.* (*22*). To be specific, DNA replication initiation is triggered when the concentration of free DnaA-ATP exceeds a critical threshold $\left[A_f^c\right]$. The free concentration is calculated as $\left[A_f^{ATP}\right] = (A^{ATP} - A_b)/V$, where $A_f^{ATP}$ is the total number of DnaA-ATP molecules, $A_b$ represents the number bound to high-affinity sites, and $V$ is cell volume. The dynamics of the total DnaA pool $(A)$ are governed by a growth-rate-dependent synthesis term that accounts for gene dosage and autoregulation: $\frac{dA}{dt} = \frac{\phi_{P0}\lambda V}{1+\left(\left[A_f^{ATP}\right]/K_D^P\right)^n}$. Meanwhile, the dynamics of the active DnaA-ATP pool $(A^{ATP})$ are described by a balance of three key processes: $\frac{dA^{ATP}}{dt} = (\alpha_l V + \alpha_{d1} N_{d1} + \alpha_{d2} N_{d2}) \frac{[A^{ADP}]}{K_D + [A^{ADP}]} - (\beta_{datA} + \beta_{rida}) N_o \frac{\left[A_f^{ATP}\right]}{K_D + \left[A_f^{ATP}\right]} + \frac{\phi_{P0}\lambda V}{1+\left(\left[A_f^{ATP}\right]/K_D^P\right)^n}$

The first term captures nucleotide exchange promoted by lipid membranes and DARS sites, the second represents inactivation via *datA* and RIDA, and the third includes newly synthesized DnaA-ATP.

This formulation relies on several key biochemical parameters that define the molecular regulation of DnaA. The total DnaA concentration in the cell is denoted by $[D_T]$, while the concentration of active DnaA-ATP is represented as $[A^{ATP}]$, and the concentration of inactive DnaA-ADP form is calculated as $[A^{ADP}] = \frac{(A - A^{ATP})}{V}$, where $V$ is the cell volume. The binding affinity of DnaA to its own promoter is governed by the dissociation constant $K_D^P$, with a cooperativity factor $n$ describing the Hill-type activation of *dnaA* gene expression. The rate of DnaA-ATP regeneration is mediated through multiple pathways: phospholipid membranes (with rate $\alpha_l$), and chromosomal DARS sites (DARS1 and DARS2, with activation rates $\alpha_{d1}$and $\alpha_{d2}$, respectively). The strength of the switching effect was further modulated by a factor $W_s = 0.5$ in the maintext. Inactivation of DnaA-ATP occurs via the *datA* locus and RIDA complex at rates $\beta_{datA}$ and $\beta_{rida}$. The overall sensitivity of these activation and inactivation reactions is modulated by a common dissociation constant $K_D$. The copy numbers of regulatory sites—$N_{d1}$ (DARS1), $N_{d2}$ (DARS2), and $N_o$ (*oriC*) — are dynamically updated during simulation based on replication progression. Additionally, $\phi_{p0}$ represents the basal ratio of DnaA relative to total cellular protein, which scales with the growth-dependent synthesis rate $\alpha_A$. The stochasticity of this model was introduced to the DnaA expression level $\phi_{p0}$ with white noise updated at each replication cycle. All parameters concerning the DnaA-ATP/ADP switching followed Berger *et al.* 2022 (*22*).

##### 2.3.2. The implementation of titration-switch model

The titration-switch model was implemented based on the initiation titration model. The update of cell stat was replaced in the main loop as follows:

**Parameters:**

λ : Cell mass growth rate
$DT = \ln(2)/\lambda$ : Doubling time
$TT = 100DT$ : Total simulation time
$dt = 0.2$ min : Time step
$nbox_{chrom} = 150$ : Number of DnaA boxes per chromosome
$C$ : Duration of DNA replication (computed from $\lambda$)
$[D_T] = 400\ \mu\text{m}^{-3}$ : Total DnaA concentration $[D_T] = 400\ \mu\text{m}^{-3}$;
$K_D^P = 400\ \mu\text{m}^{-3}$ : Promoter dissociation constant;
$\phi_{p0} \in 0.002\%\ to\ 0.1\%$ : The DnaA protein ratio relative to all proteins;
$n = 5$ : The cooperativity of *dnaA* expression;
$\alpha_l = 750\ \mu\text{m}^{-3}\text{h}^{-1}$ : The activation rate of DnaA-ATP by lipid;
$\alpha_{d1} = 100\ \mu\text{m}^{-3}\text{h}^{-1}$ : The activation rate of DnaA-ATP by the DARS1 site;
$\alpha_{d2} = 643\ \mu\text{m}^{-3}\text{h}^{-1}$ : The activation rate of DnaA-ATP by the DARS2 sites;
$\beta_{datA} = 600\ \mu\text{m}^{-3}\text{h}^{-1}$ : The deactivation rate of DnaA-ATP by *datA*;
$\beta_{rida} = 500\ \mu\text{m}^{-3}\text{h}^{-1}$ : The deactivation rate of DnaA-ATP by RIDA;
$K_D = 50\ \mu\text{m}^{-3}$ : The dissociation constant of DnaA activation;
$P_{d1} = 0.1 \cdot C$ : Location of DARS1 on chromosome;
$P_{d2} = 0.2 \cdot C$ : Location of DARS2 on chromosome;
$[A_f^c] = 200\ \mu\text{m}^{-3}$ : The initiation threshold of free DnaA-ATP;
$\epsilon_A = 0.15$ : The noise strength on the DnaA expression level
$W_s = 0.5$ : Strength factor for the switching effects
$AboxRange = 0.8$ : Distribution range of DnaA boxes on chromosome

**While $t < t_{max}$ do**

➢ Update cell state:

### Cell mass growth and DnaA synthesis

$V = V + \lambda \cdot V \cdot dt$

$A = A + \frac{\phi_{P0}\lambda V}{1+\left(\left[A_f^{ATP}\right]/K_D^P\right)^n} \cdot dt$

$A^{ATP} = A^{ATP} + W_s(\alpha_l V + \alpha_{d1}N_{d1} + \alpha_{d2}N_{d2})\frac{[A^{ADP}]}{K_D+[A^{ADP}]} - W_s(\beta_{datA} + \beta_{rida})\, N_o \frac{\left[A_f^{ATP}\right]}{K_D+\left[A_f^{ATP}\right]} + \frac{\phi_{P0}\lambda V}{1+\left(\left[A_f^{ATP}\right]/K_D^P\right)^n} dt$

$A^{ADP} = A - A^{ADP}$

### Get gene amounts and number of DnaA-boxes

$DNA = sum(DNAcopy)$

$N_{d1} = DNAcopy(P_{d1})$

$N_{d2} = DNAcopy(P_{d2})$

$OriC = DNAcopy(1)$

$Abox = DNA(1{:}AboxRange) \cdot nbox_{chrom}$

➢ DNA replication initiation:

**If** $(A^{ATP} - Abox)/V > [A_f^c]$

### Start a new replication fork at origin of the DNAcopy vector and double it

$nrep = nrep + 1$

$DNAcopy(1) = 2 \cdot DNAcopy(1)$

$Prep(nrep) = 1$

### Reset stochastic DnaA synthesis rate:

Draw a random unit Gaussian variable $\eta$,

$\alpha = \alpha_0 + \epsilon_A \eta$

### Record cell volume, initiation time and origin number at initiation

$V_i(nrep) = V/OriC$

$T_i(nrep) = t$

$O_i(nrep) = OriC$

**end**

➢ Update DNAcopy:

**end**

##### 2.3.3. Effects of the switching strength

By simulating this model, we successfully captured the fundamental quantitative relationships observed in bacterial cell cycle control, specifically the II adder correlations (Fig. S16) with parameters presented below in the pseudo code. The change of expression level is now applied by $\phi_{P0}$ has the same effect as in changing DnaA synthesis rate $\alpha$ in the classic titration model.

We then systematically tested the effect of DnaA's switching strength (applied on all the active rates $\alpha_l$, $\alpha_{d1}$, $\alpha_{d2}$ and deactivate rates $\beta_{datA}$, $\beta_{rida}$). By increasing this switching strength factor, we observed a direct correlation with the II adder slope, particularly in fast growth conditions. As shown in Fig. S17, increasing the switching strength caused the II adder slope to approach 0.

##### 2.3.4. The extrusion model

The classic DnaA titration model fails to predict the persistence of replication initiation cycles even after DnaA synthesis is abruptly halted. The extrusion model was proposed to resolve this discrepancy by extending the titration framework by introducing a hypothetical extruder protein, which actively displaces DnaA from chromosomal DnaA-boxes. This mechanism provides a dynamic release of bound DnaA into the free pool, effectively acting as a "molecular extruder" that senses deviations between biomass growth and DNA content. Unlike passive titration, where DnaA availability depends solely on synthesis and origin duplication, this model allows for reactivation of DnaA through competitive displacement, enabling sustained oscillations in free DnaA concentration and continued initiation events even in the absence of new DnaA synthesis (*15*).

The extrusion model retains the core structure of the classic titration model—DnaA synthesis, binding to DnaA-boxes, and threshold-based initiation—but introduces two critical innovations: (1) a novel regulatory component: an extruder protein ($H$), hypothesized to actively displace DnaA from its chromosomal binding sites. This extruder is constitutively expressed at a constant rate per unit volume, independent of the replication cycle or DnaA regulation, leading to continuous accumulation proportional to biomass growth: $dH/dt = \alpha_H V$ with synthesis rate $\alpha_H = 100\ \mu m^{-3} h^{-1}$ . (2) the revised calculation of free DnaA concentration, which now accounts for the competitive action of the extruder: $[A_f] = (A - (A_b - H))/V$, under condition of $(A_b - H) \geq 0$ and $(A - (A_b - H)) \geq 0$. The extruder concentration played similar role as the DnaA expression level which changed primarily the average initiation mass. Extruder expression level has the same effect as *dnaA* expression level (*15*).

To simulate this model, one can change the pseudo code based on titration model as follows:

**Parameters**

$\alpha_H = 100$ : Extruder synthesis rate in $\mu m^{-3} h^{-1}$

**While $t < t_{max}$ do**

- Update cell state:

  \# Extruder synthesis

  $H = H + \alpha_H \cdot V \cdot dt$

  \# Get gene number of active DnaA-boxes to bind DnaA protein

  $Abox = N_{datA} \cdot nbox_{datA} + DNA(1{:}AboxRange) \cdot nbox_{chrom} - H$

  If $Abox < 0$

  $Abox = 0;$

  end

- DNA replication initiation:
- Update DNAcopy:

**end**

#### 2.4. The IDC model

##### 2.4.1. Model construction

Based on the ID adder principle uncovered in this study, we formulate cell division as occurring when the cell volume reaches a target mass that is set at the time of replication initiation. Specifically, the division volume $V_d$ is modeled as:

$$V_d = V_i + \Delta V_{id} \times oriC$$

where $V_i$ is the initiation volume, $\Delta V_{id}$ is a stochastically varying volume increment, and $oriC$ is the number of origins at initiation. In our simulation framework, whenever the current cell volume $V$ reaches or exceeds any of the precomputed division mass thresholds (i.e., the $V_d$ values were set at prior initiations), a division event is triggered.

To capture inter-cycle correlations (ICC) in the added volume, we describe the dynamics of $\Delta V_{id}$ using a stochastic Ornstein–Uhlenbeck (OU) process:

$$\mathrm{d}\Delta V_{id} = \kappa(\Delta V_{id} - \overline{\Delta V_{id}}) + \epsilon_{id}\eta$$

where the average ID added volume per origin $\overline{\Delta V_{id}} = \overline{V_d} - \overline{V_i}$ is derived from the experimentally measured initiation mass and division mass. $\overline{V_d}$ is estimated using the relation from (*7*): $\overline{V_d} = m_0/ln(2)(0.28 + 0.99\lambda)$ with $m_0$=1 representing the unit cell mass and $\lambda$ the growth rate (Fig. S18). The term $\epsilon_{id}\eta$ introduces Gaussian white noise accounting for intrinsic stochasticity in volume addition. The parameter $\kappa$ quantifies the strength of inter-cycle correlation in $\Delta V_{id}$, its value is determined by fitting to the empirical ICC curve shown in Figure 2E of the main text. Here, we adopt the OU model not because it reflects a known mechanistic basis for $\Delta V_{id}$ regulation, but because it provides the simplest mathematically tractable framework capable of incorporating a tunable degree of memory (i.e., ICC) into the stochastic dynamics of the added volume.

##### 2.4.2. Model implementation

To incorporate the inter-cycle correlation (ICC) and the ID adder principle into our framework, we build upon the simplest classic titration model and extend it with the following key features of cell division:

- **Initiation-to-Division coupling**: Each initiation event establishes a future division target through a dynamically computed $V_d$, implementing a tunable initiation-initiation adder strategy that links replication onset to subsequent cell division.
- **Stochasticity with autocorrelation**: Biological variability is introduced by adding noise to volume increment added per origin ($\Delta V_{id}$) at every initiation, thereby generating natural cell-to-cell heterogeneity across generations. The parameter $\kappa$ enables modeling of heritable fluctuations in ID added mass with autocorrelation. The autocorrelation effect between successive cell cycles was implemented via an Ornstein–Uhlenbeck–type process with $\Delta V_{id} = -\kappa(\Delta V_{id} - V_{id}^0) + \epsilon_{id} V_{id}^0 \eta$, where $\eta$ is a Gaussian noise , $\epsilon_{id}$ is the noise level and $V_{id}^0$ is the average added mass.
- **Symmetric division**: Upon reaching its assigned $V_d$ , the cell divides instantaneously. The full simulation implicitly enforces symmetric division: cellular

volume and molecular counts—including the DNA structure vector—are halved between daughter cells. At division, the ploidy count is doubled (reflecting completed replication), the division time and mass ($V_d$) are recorded, and the corresponding $V_d$ threshold is set to infinity to ensure it is not reused—thus guaranteeing that each initiation sets exactly one division event, in line with the biological logic of the ID adder.

To simulate this model, one can change the following aspect based on titration model as follows:

**Parameters**

$\Delta V_{id}^0 = V_D^0 - V_i^0$ : Average added mass between replication initiation and cell division

$\kappa$ : Autocorrelation between $\Delta V_{id}$, fitted from experiment data

$\epsilon_{id} = 0.15$ : Noise level on $\Delta V_{id}$

**While $t < t_{max}$ do**

- Update cell state:
- Check for initiation of DNA replication:

  **If** $(A - Abox)/V \ > \ [A_f^c]$

  # Start a new replication fork at origin of the DNAcopy vector and double it
  # Reset stochastic division target
  Draw a random unit Gaussian variable $\eta$,
  $\Delta V_{id}(nrep) = -\kappa(\ \Delta V_{id}(nrep-1) - \Delta V_{id}^0) \ + \ \Delta\overline{V_{id}}\epsilon_{id}$

  $V_d^{target}(nrep) = V_i(nrep) + \Delta V_{id}(nrep) \cdot OriC$

  # Record cell volume, initiation time and origin number at initiation
  **end**
- Update active replication forks
- Check if any division targets are reached

  **If** $V \geq V_d^{target}$:

  $nd \ = \ nd + 1$
  # Record division mass, division time and cell number
  $T_d(nd) \ = \ t$
  $N(nd) \ = \ 2N(nd-1)$
  $V_d(nd) \ = \ V/N$

  **end**

**end**

##### 2.4.3. Simulation results

To investigate how the IDC (Initiation-to-Division Correlation) model behaves under diverse physiological conditions, we conducted stochastic simulations across two distinct growth rates, corresponding to growth rates of 0.3 $h^{-1}$ (slow) and 1.8 $h^{-1}$ (fast). The results are summarized in Fig. S19, which combines time-resolved trajectories of cell mass with intergenerational correlations in division behavior, offering a comprehensive assessment of the model's predictive power.

Through scatter plots of rescaled division mass ($V_d$), added mass ($\Delta V_d$) and elapsed time ($\tau_{dd}$) versus rescaled division mass ($V_d$) at two different growth rates (Fig. S19CD), we see that $\Delta V_d$ is uncorrelated with $V_d$ at fast growth rate (Fig. S19D middle panel), confirming the DD adder phenomenon while the other correlations deviates from either sizer nor timer principles (non-zero slopes). These results show a qualitative agreement with the experimental measured DD correlations (Figure S2).

In addition to the well-established correlations among inter-initiation (II), initiation-to-division (ID), and division-to-division (DD) intervals, we further tested the predictive power of the Initiation-to-Division Coordination (IDC) model by analyzing birth-to-initiation relationships. Specifically, we quantified the slope of rescaled initiation mass versus rescaled birth mass using linear regression and compared the experimentally measured slopes with IDC model predictions across a range of growth conditions (Fig. S20). Both the model and experimental data show a consistent decrease in this slope with increasing growth rate, indicative of a progressive loss of birth-mass memory at replication initiation.

##### 2.4.4. Noise effects on the cell division

The introduction of noise plays a pivotal role in stochastic simulations of cellular processes, as it captures the inherent randomness present in biological systems. In a growing cell, two primary sources contribute to this stochasticity: fluctuations in cell mass (or volume) growth and variability in gene expression. While both are essential for realistic modeling, they exert distinct influences on cell cycle regulation.

Specifically, noise associated with cell growth—arising from variations in nutrient uptake, metabolic activity, or biosynthetic rates—does not significantly affect the correlations observed at the time of replication initiation. However, this same growth noise has a pronounced impact on division-to-division correlations. Because division is often coupled to mass or added mass, any variability in growth dynamics propagates into the timing and mass at division, thereby shaping intergenerational division correlations (Fig. S21).

##### 2.4.5. Comparison to other division models

In recent years, efforts to explain the robust adder correlations observed in bacterial cell mass control have led to the development of several distinct theoretical frameworks. Among the most prominent are the Independent Double Adder (IDA) model, the Replication Double Adder (RDA model), and the Concurrent Cell-Cycle Processes model (CCCP model). These models diverge in their underlying principles for division timing control (*23-27*). While commonalities in reproducing DD and II adder phenomena exist among these models, the causal mechanisms governing bacterial division timing continue to elude definitive resolution. Considering the IDA, RDA, and CCCP models, all of which are initially rooted in the constraints of the II and DD adder phenomena, with a primary focus on unraveling the mechanisms governing division timing control.

The IDA model appears inconsistent with our observations. It treats initiation and division as independent adder processes relative to birth (*25*), which fails to capture the origin-scaling effects we observe under initiation perturbations. In both cases, the predicted responses to altered initiation mass do not match the resilience and scaling behavior seen in our experiments.

The initiation mass perturbation experiments presented in this study lend support to the ID adder concepts proposed by the RDA model (*26, 27*) but introduces ICC to correct its growth rate dependent correlations. Our experimental perturbations of initiation mass provide critical leverage for discriminating among these competing frameworks. The results align most closely with predictions of the RDA model, which posits that both replication initiation and cell division are governed by adder-like processes tied to the number of origins: two fixed volumes per origin are added, one between successive initiations and another—between initiation and division. This dual-adder structure naturally accounts for the persistence of adder behavior even when initiation timing is experimentally shifted—a feature our data support.

The CCCP model presents a more flexible alternative. As a modular framework that allows multiple cell-cycle events—such as replication -initiation, -termination and cell division—to proceed concurrently under shared or distinct timers, it can be tuned to accommodate a wide range of empirical patterns (*28, 29*). Indeed, by incorporating growth-rate–dependent parameters, the CCCP model can also reproduce the recently uncovered modulation of adder strength with growth conditions—a phenomenon that places stringent constraints on any viable theory of mass control. Our IDC model is not unique in this regard; it joins the CCCP framework as one of several plausible architectures capable of explaining both the canonical adder correlations and their nuanced dependence on physiological context.

We notice that under extreme perturbations—for example, those that greatly prolong $D$ by further reducing the expression level of division-related genes (Figure S8)—the ID adder phenomenon can break down. In such extreme cases, it is plausible to combine the IDC model with a DD adder–determined division process through a logical AND gate, forming a concurrent-type model to represent the full regulation of cell division.

To incorporate this biological complexity, we expanded our IDC model by integrating it with a division process governed by a DD adder mechanism, using a logical AND gate to couple the two processes (IDC-DD concurrent model). This extension allows us to simulate $D$-perturbations experimentally, such as those induced by the A22 antibiotic (*28*), by modulating the average added mass associated with the DD adder. Utilizing this enhanced IDC-DD concurrent model, we were able to reproduce trends consistent with those reported by Colin et al., specifically regarding the dependence of the ID adder slope on the $D$ period and the relationship between the ID and DD adder slopes

(Fig. S22). These findings suggest that the IDC framework can effectively account for *D*-dependent division control without necessitating an ID timer mechanism.

Thus, while our findings favor mechanisms that explicitly link division to the number of active origins—as embodied in the RDA model—they do not rule out adaptable frameworks like CCCP. Instead, they underscore the need for models that integrate molecular regulation (e.g., DnaA dynamics and *OriC* licensing) with system-level mass control, and that remains responsive to environmental modulation. Our IDC model offers one such synthesis, based in biochemistry yet capable of reproducing emergent population-level regularities. The comparison of our model and existing models is summarized in Table S4.

##### 2.4.6. About the causal correlation

In a recent study, by utilizing conditional independent tests, Kar *et al.* suggested that chromosome replication-initiation exclusively governs cell division in slow-growth conditions, whereas additional factors play during fast-growth conditions (*30*). Numerical simulation of the titration-correlation model yields results concordant with their observations and suggests that the ICC of $\Delta_{id}$ may act as an “additional factor” hypothesized by Kar *et al.* (*30*) (Fig. S23).

#### 3. Supplementary Figures and Legends

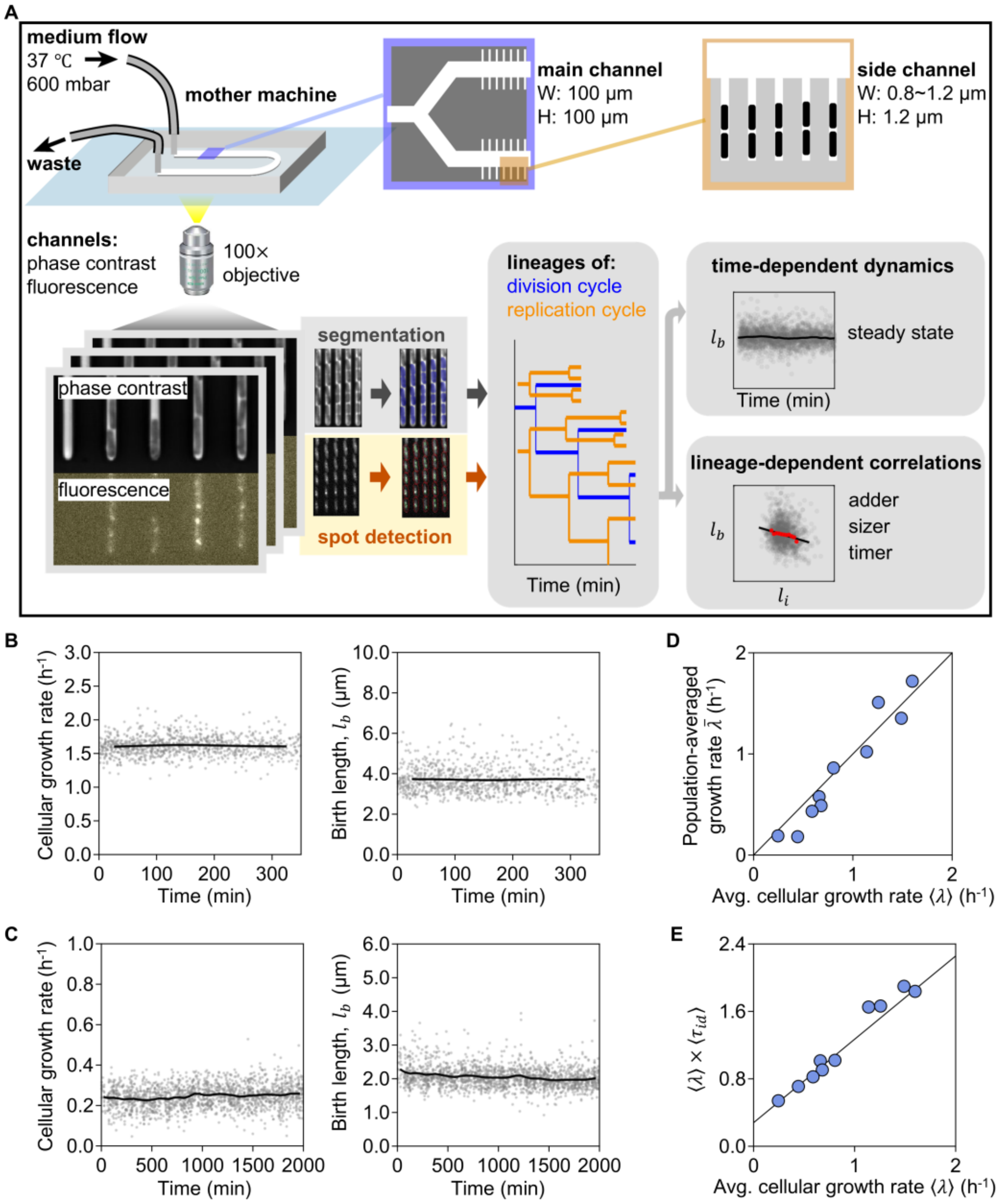


**Figure S1. Single-cell experimental system and validation of steady-state growth.** (**A**) Workflow for single-cell analysis using mother machine microfluidics with continuous media perfusion. Time-lapse microscopy in phase contrast and fluorescence channels enabled lineage tracking and quantification of all cell-cycle parameters. (**B**, **C**) Temporal stability of single-cell growth rates and birth sizes in (**B**) RDM+glucose and (**C**) MOPS+alanine. Gray points, single-cell data; black line, population mean over time. (**D**) Correlation between the average cellular growth rate of our strain (ZH101) and reported population-averaged values for MG1655 under the same media *(4)*. The black line indicates $y = x$. (**E**) The product of the average time from initiation to division ($\langle \tau_{id} \rangle$) and $\langle \lambda \rangle$ as a function of averaged cellular growth rate ($\langle \lambda \rangle$). The black line shows the previously established population-level relationship *(4)*, demonstrating that our single-cell data recapitulates known physiology.

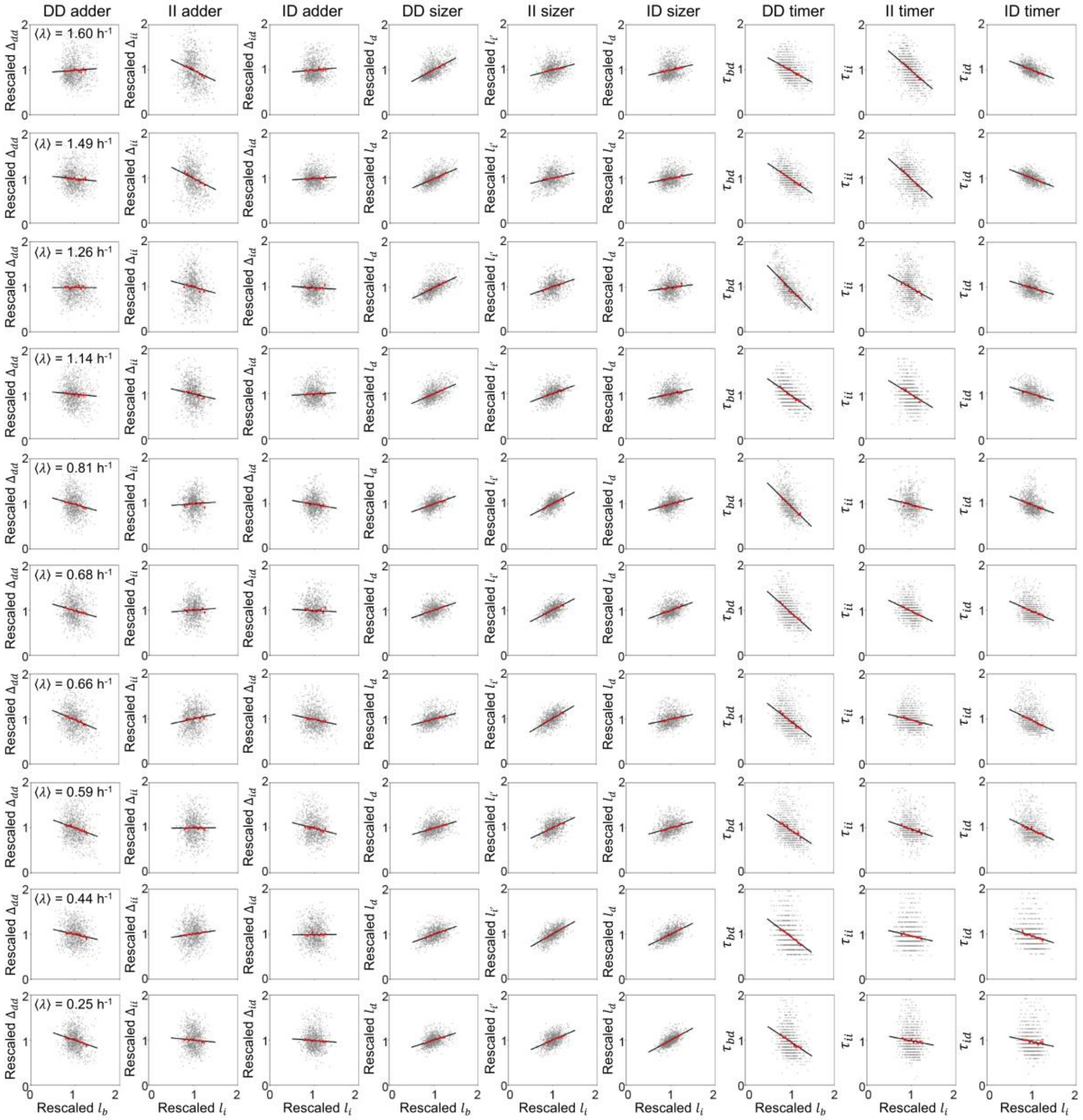


**Figure S2. Comprehensive single-cell correlations across ten growth media.** Scatter plots for nine cell-cycle correlations across all ten growth conditions, testing adder, sizer, and timer models for the DD, II, and ID intervals. Parameters are defined as: $l_b$, $l_d$, $l_i$, $l_{i'}$, size at birth, division, current initiation, and next initiation, respectively; $\Delta$ and $\tau$ represent size added and time elapsed for DD, II, and ID intervals. All parameters were normalized by their condition-specific mean. Data are presented as single-cell points (gray), binned averages ± SEM (red), and a linear fit to all single-cell data without outliers (black line). $n \geq 1000$ cells per condition. See appendix file 1 for media details and exact $n$.

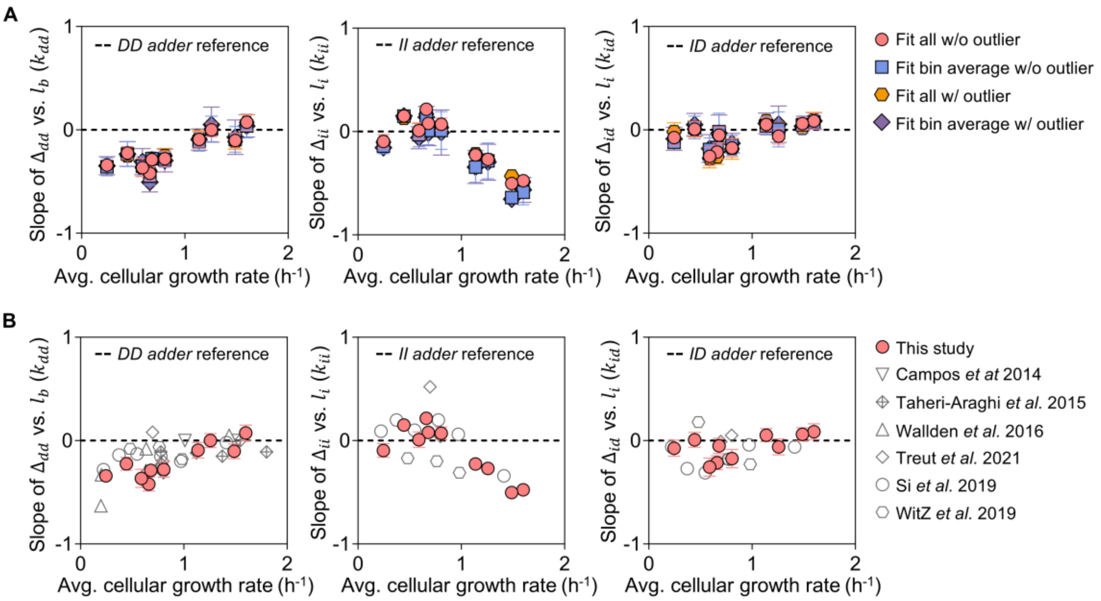


**Figure S3. Robustness of adder correlations to analytical methods and consistency with prior studies.** (**A**) Growth-rate dependence of the adder slopes $k_{dd}$ (left), $k_{ii}$ (middle), and $k_{id}$ (right) calculated using four distinct methods. Error bars, 95% confidence intervals. The dashed line indicates the expectation ($k = 0$) for a perfect adder. (**B**) Comparison of adder slopes from this study with those re-analyzed from published datasets *(1, 6, 14, 17, 19, 25)*. The dashed line indicates $k = 0$. Datasets with $n < 300$ were excluded to ensure statistical robustness.

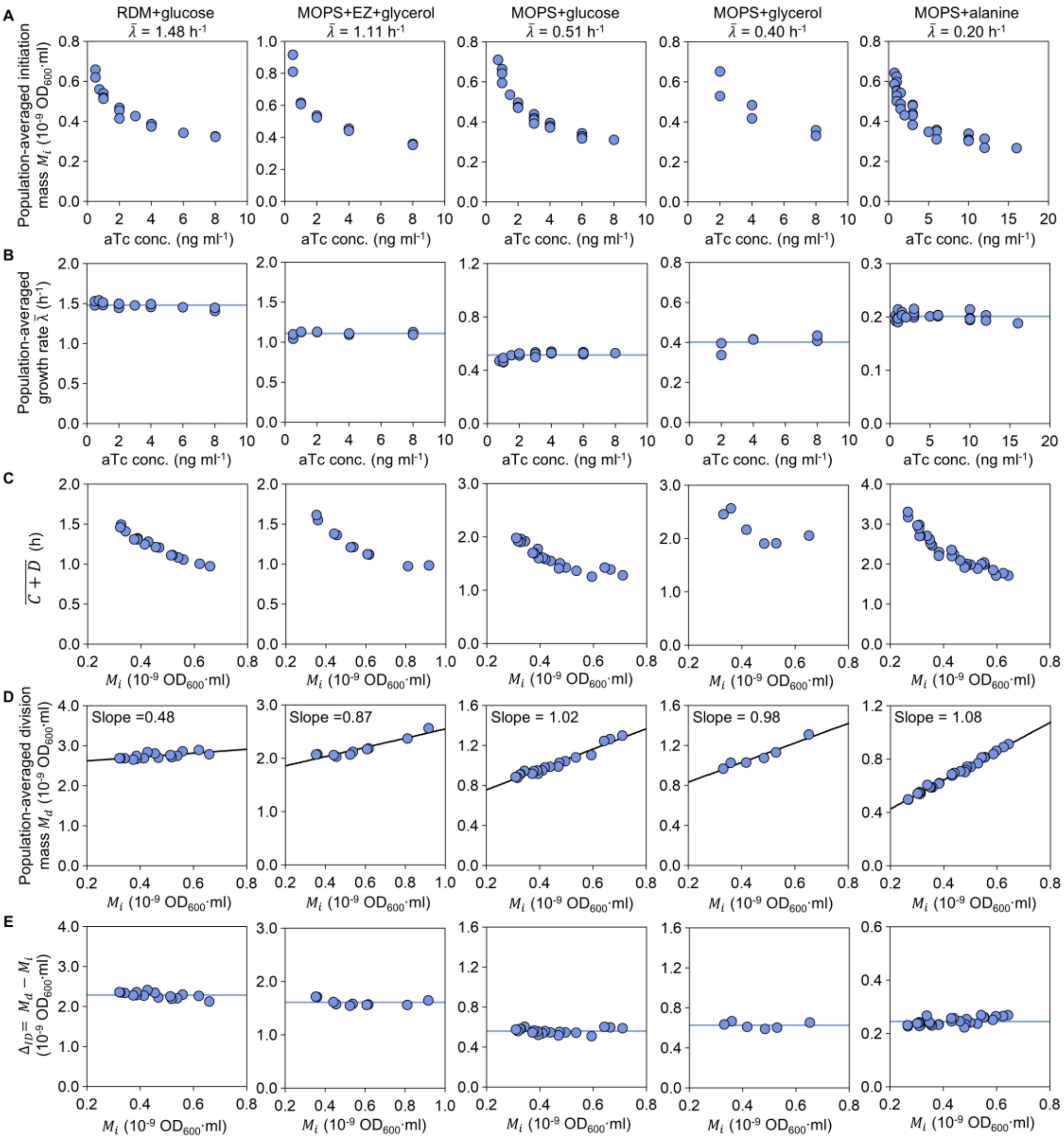


**Figure S4. The ID adder principle under DnaA titration across growth conditions.** Population-level analysis of the DnaA-titratable strain (CLL1) under various growth media. (**A**) Average initiation mass ($M_i$) as a function of aTc concentration. (**B**) Average growth rate ($\bar{\lambda}$) as a function of aTc concentration. The blue line indicates the mean growth rate for each condition. (**C**) Average initiation-to-division time as a function of initiation mass ($M_i$). (**D**) Average division mass ($M_d$) as a function of initiation mass ($M_i$). The black line is a linear fit (slope indicated). (**E**) The ID added mass ($\Delta_{ID}$) remains constant across the titrated range of $M_i$. The blue line indicates the mean $\Delta_{ID}$. Data points are from ≥2 independent experiments.

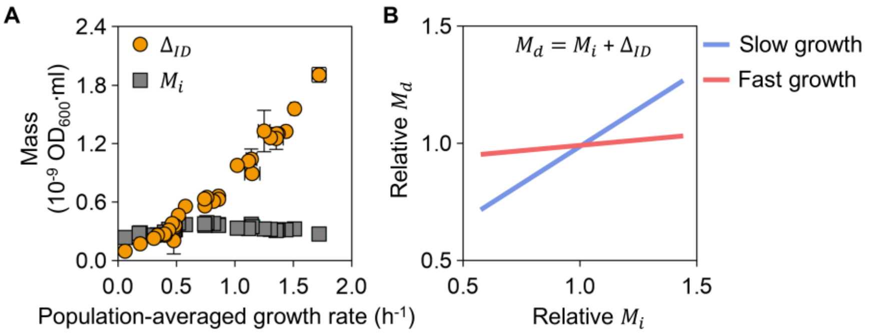


**Figure S5. The effect of added mass magnitude on the initiation-division relationship.** (**A**) Published data *(4)* for MG1655 showing the population-averaged ID added mass ($\Delta_{ID}$) and initiation mass ($M_i$) as functions of growth rate. (**B**) Conceptual model: The slope of the $M_d$ vs. $M_i$ relationship is governed by the relative magnitude of $\Delta_{ID}$ and $M_i$. At slow growth ($\Delta_{ID} \approx M_i$), the slope is ~1; at fast growth ($\Delta_{ID} \gg M_i$), the slope becomes shallow, as mathematically required by the ID adder principle ($M_d = M_i + \Delta_{ID}$).

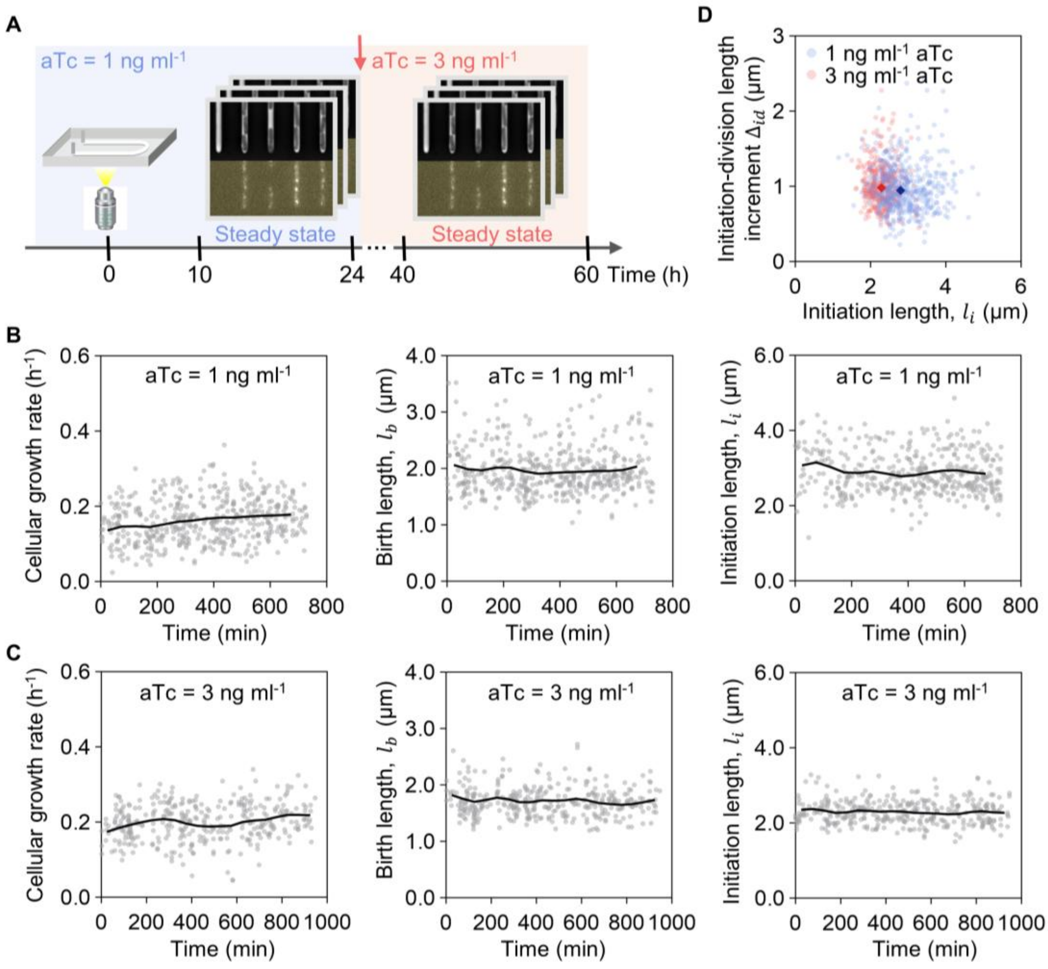


**Figure S6. Single-cell analysis of the DnaA-titration strain.** (**A**) Workflow for single-cell analysis of the DnaA-titration strain, using mother machine microfluidics with continuous media perfusion and inducer shifts (see Supplementary Materials). (**B**, **C**) Temporal stability of single-cell growth rates, birth size ($l_b$) and initiation size ($l_i$) for CLL1 strains with DnaN-Ypet cultured in MOPS + alanine supplemented with (**B**) 1 ng ml$^{-1}$ aTc and **(C)** 3 ng ml$^{-1}$ aTc. Gray points, single-cell data; black line, population mean over time. (**D**) Initiation-to-division length increment ($\Delta_{id}$) versus initiation length ($l_i$). Colored points, single-cell data; colored diamonds: the corresponding means. n = 502 (1 ng ml$^{-1}$ aTc) and n = 386 (3 ng ml$^{-1}$ aTc) cells.

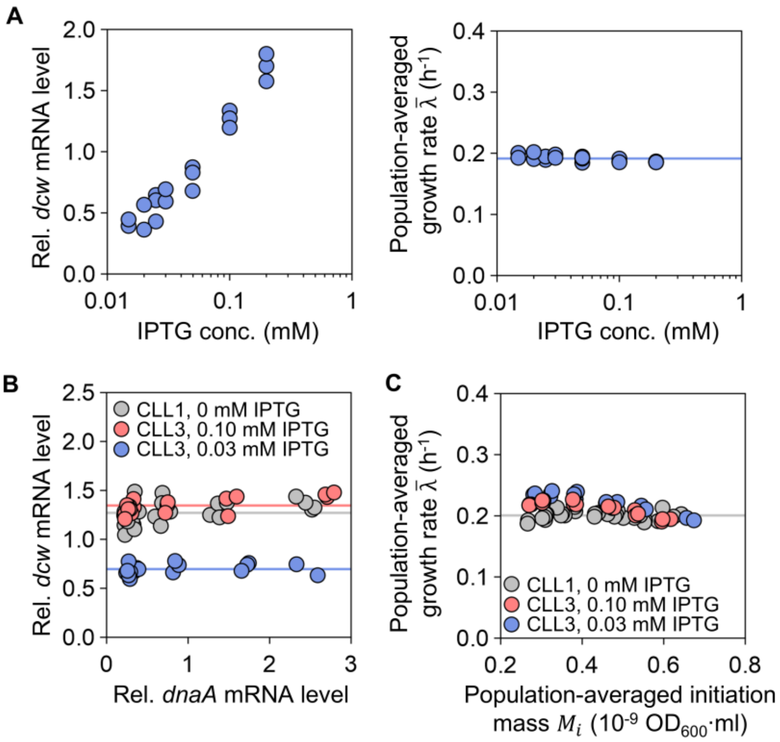


**Figure S7. Orthogonal control of division and initiation in titration strains.** (**A**) Left: Relative *dcw* mRNA level in the *dcw*-titratable strain (CLL2) as a function of IPTG concentration. Right: Growth rate ($\bar{\lambda}$) is unaffected by IPTG titration. The blue line indicates the mean. (**B**) In the dual-control strain (CLL3), titrating *dnaA* expression (via aTc) does not affect *dcw* expression (via fixed IPTG), confirming orthogonal control. Colored lines indicate mean *dcw* mRNA levels for different strain/induction conditions. (**C**) Growth rate ($\bar{\lambda}$) remains invariant with initiation mass ($M_i$) in both the CLL1 (*dnaA*-titratable) and CLL3 (dual-titratable) strains. Gray lines indicate the mean growth rate for CLL1. Data points are from ≥3 independent experiments.

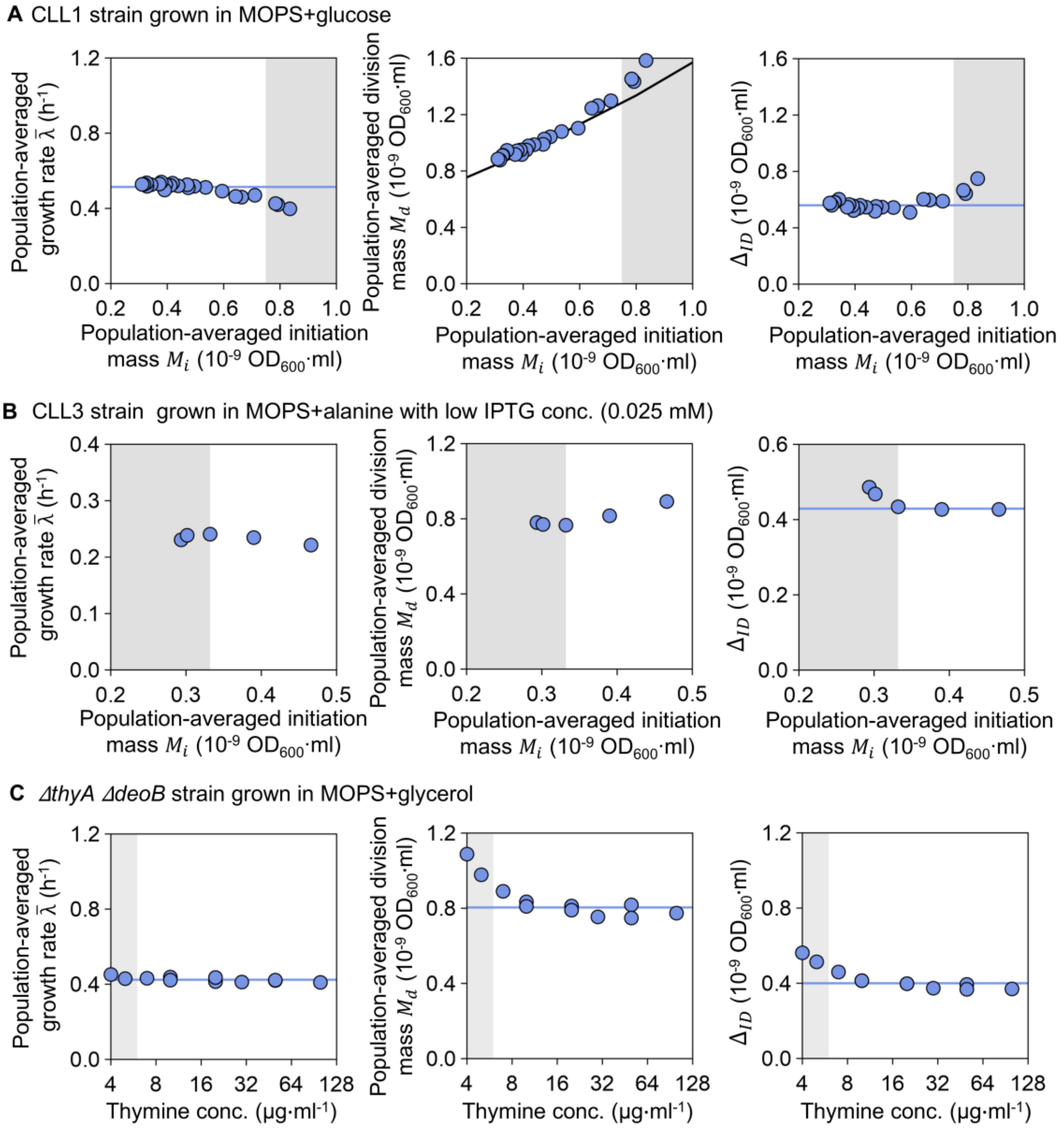


**Figure S8. Breakdown of the ID adder principle under extreme perturbations.** (**A**) In the *dnaA*-titratable strain (CLL1) under MOPS+glucose, severe aTc concentration reduction leads to excessively large initiation mass, causing a decline in growth rate and the collapse of the ID adder (right shaded region). (**B**) In the dual-control strain (CLL3) under severe division limitation (0.025 mM IPTG), division becomes uncoupled from initiation at small $M_i$ (left shaded region). Here, division mass is fixed by the limited division machinery, and the added mass ($\Delta_{ID}$) decreases with increasing initiation mass, violating the ID adder. (**C**) In the *thyA deoB* double knockout strain, the division mass ($M_d$) and added mass ($\Delta_{ID}$) increases with decreasing thymine concentrations.

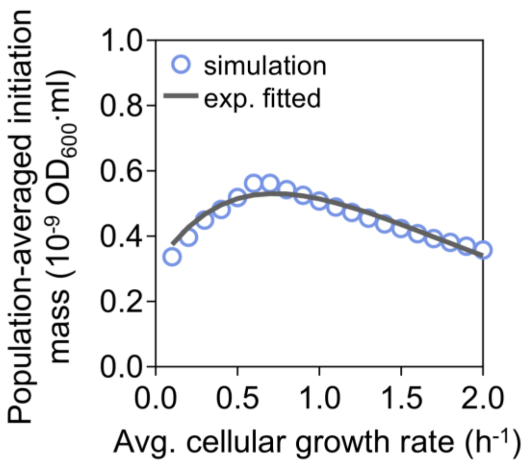


**Figure S9. Parameter fitting for DnaA synthesis rate ($\alpha$).** For the simulated growth rates varies from $0.1\ \mathrm{h}^{-1}$ to $2\ \mathrm{h}^{-1}$ (blue circles), $\alpha$ was chosen as shown in Table S3, to mimic the experimentally fitted average initiation mass over growth rate ($V_i = \frac{m_0}{\ln 2}(0.28 + 0.99\lambda)e^{-(0.28+0.99\lambda)}$ (*7*), dark gray line).

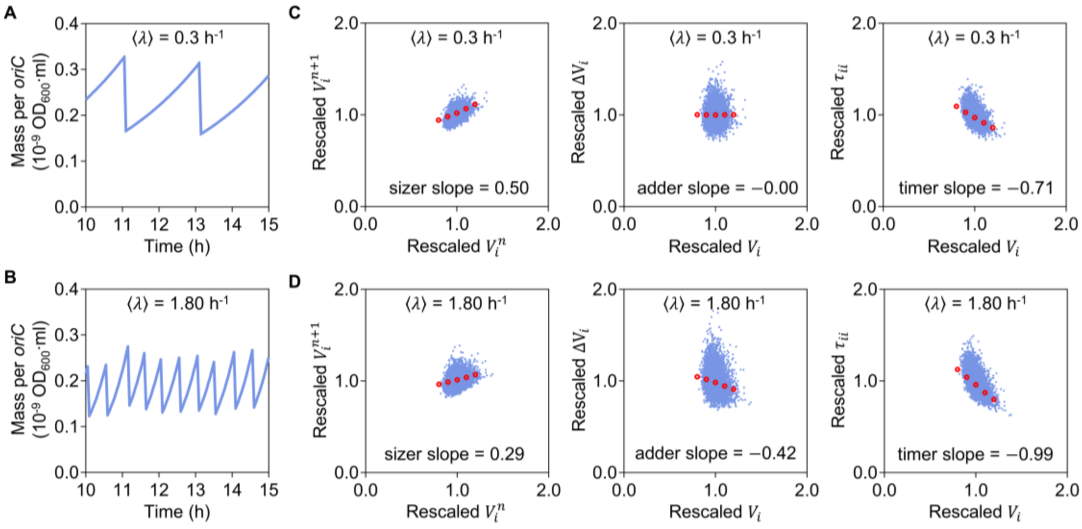


**Figure S10. Predicted mass per *oriC*: dynamics and single-cell correlations between successive initiations from the stochastic titration model. (AB)** Time trajectories of cell mass per *oriC* over 5 hours for two distinct averaged cellular growth rates ($\langle\lambda\rangle = 0.3$ h$^{-1}$ and $\langle\lambda\rangle = 1.8$ h$^{-1}$), showing oscillations in cell mass per *oriC* over time. **(CD)** Scatter plots for sizer, adder, and timer correlations: Sizer, rescaled initiation mass of the next round ($V_i^{n+1}$) versus current rescaled initiation mass ($V_i^n$); adder, rescaled initiation mass ($V_i$) versus rescaled increment during sequential initiation events ($\Delta V_i$); timer, rescaled time increment during sequential initiations ($\tau_{ii}$) versus rescaled initiation mass ($V_i$). Red dots represent the average trend of the scatter plots. Sizer slope, adder slope and timer slope are obtained from linear fits of the corresponding scatter data.

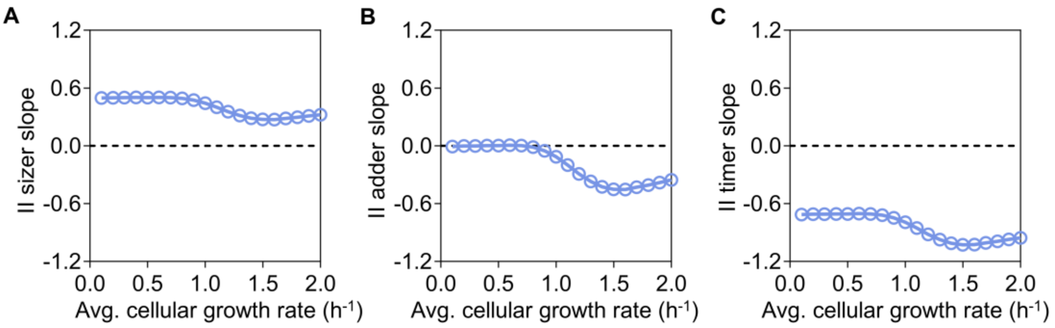


**Figure S11. Growth-rate-dependent single-cell correlations between successive initiations.** Model simulations yielded growth-rate-dependent II sizer slope **(A)**, II adder slope **(B)**, and II timer slope **(C)**. Blue circles connected by solid lines represent simulation results across growth conditions, plotted alongside the reference slopes corresponding to ideal sizer, adder, and timer behaviors (slope = 0; black dashed lines).

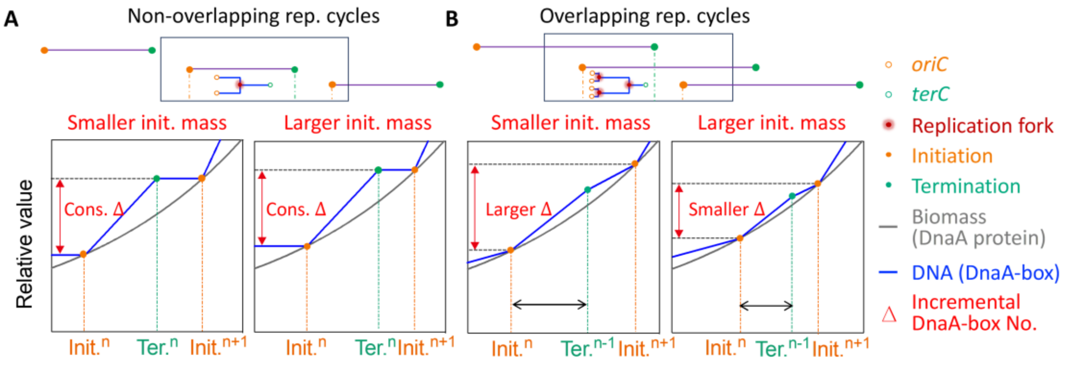


**Figure S12. Graphic illustration of the growth-rate dependence of II adder behavior. (A)** The inter-initiation time is longer than the time required for replicating the whole chromosome, current round of DNA replication terminates (Term.$^{n}$) before next round of DNA replication initiation (Init.$^{n+1}$). The incremental DnaA-box No. between consecutive replication initiation event (Δ) is defined by the DnaA-box No. per chromosome, that is independent of the cell mass at the time of DNA replication initiation. Conversely, during overlapping replication cycles **(B)**, Init.$^{n+1}$ occurs before Term.$^{n}$. The constancy of Δ is relinquished, replaced by a negative correlation with the initiation mass for Init.$^{n}$. For the display of chromosome structures under different parameters, it is recommended to use the online Cell Cycle Simulation program provided in: http://www.isynbio.org.cn/en/culture/physiology

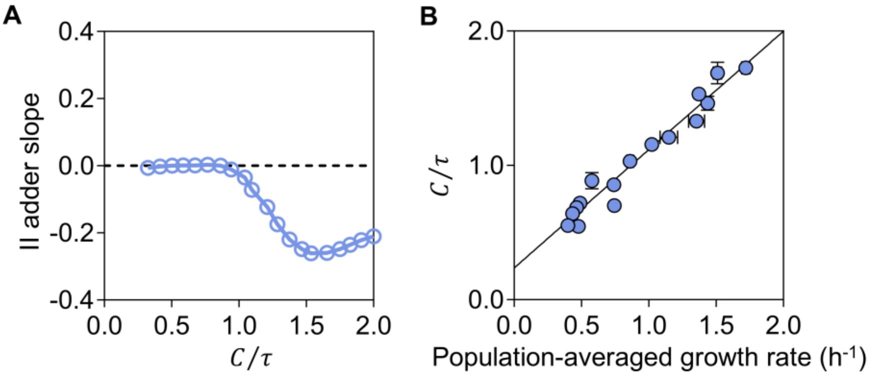


**Figure S13. Effect of the overlapping strength $C/\tau$.** (**A**) The II adder slope as a function of replication overlapping strength. Prediction derived from numerical simulation of the initiator titration model. (**B**) The growth-rate dependency of population-averaged $C/\tau$. Data points are taken from our previous study (*7*) and the solid line is a linear fit.

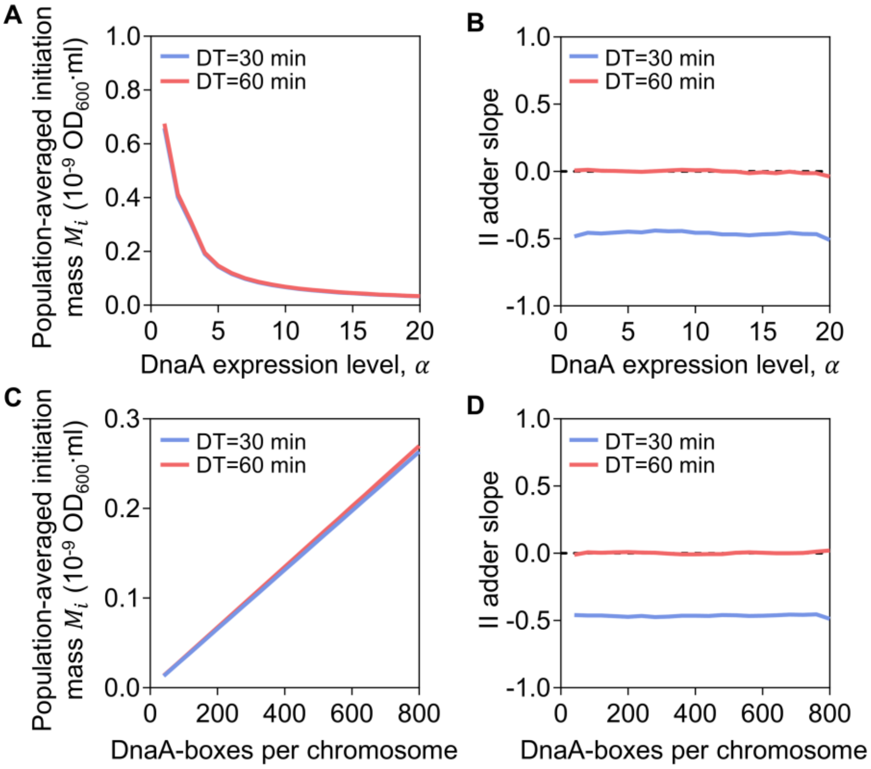


**Figure S14. DnaA synthesis rate and the number of DnaA-boxes per chromosome influence average initiation mass but not the II adder slope.** Modulating the DnaA synthesis rate (α) reduces the average initiation mass (**A**) but leaves the II adder slope unchanged (**B**) under both slow (red line) and fast (blue line) growth conditions. Increasing the number of DnaA-boxes raises the average initiation mass (**C**), yet has no effect on the II adder slope (**D**) under either slow (red line) or fast (blue line) growth condition.

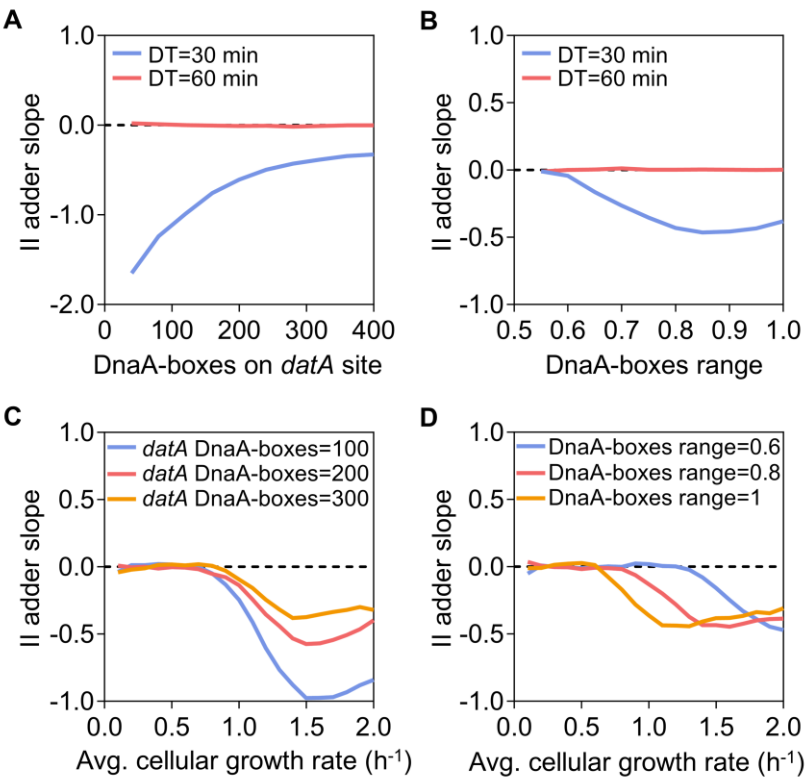


**Figure S15. The influence of DnaA-boxes number and distribution on the II adder slope under varying growth rates.** (**AB**) show how the number of DnaA-boxes at the *datA* site (parameter $nbox_{datA}$) and their chromosomal distribution range (parameter $AboxRange$) affect the II adder slope. Results are shown for fast (DT = 30 min) and slow (DT = 60 min) growth. The dashed line marks ideal adder behavior (slope = 0). (**CD**) illustrate the dependence of the adder slope on growth rate for different numbers of DnaA-boxes at *datA* site (100, 200, 300) and varying DnaA-box distribution ranges (0.6, 0.8, 1.0). Both the quantity and spatial spread of DnaA-boxes influence mass control, especially during rapid growth.

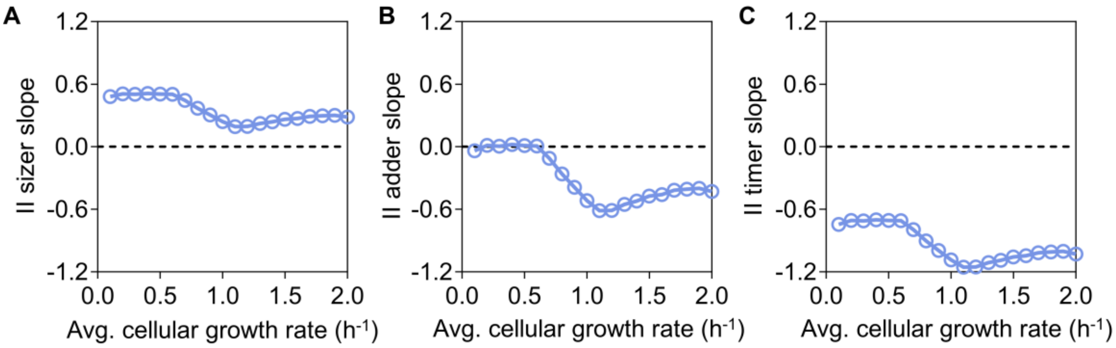


**Figure S16. Predicted growth-rate-dependent single-cell correlations between successive initiations from the titration switch model.** Model simulations yielded growth-rate-dependent II sizer slope (**A**), II adder slope (**B**), and II timer slope (**C**). Blue circles connected by solid lines represent simulation results across growth conditions, plotted alongside the reference slopes corresponding to ideal sizer, adder, and timer behaviors (slope = 0; black dashed lines).

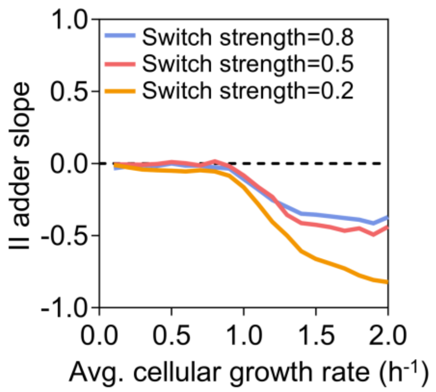


**Figure S17. Effect of switching strength on the growth-rate dependence of II adder behavior in the titration switch model.** The DNA replication initiation adder slope is plotted as a function of growth rate ($h^{-1}$) for three values of the switching strength, $W_s = (0.2, 0.5, 0.8)$. Here, the switching strength scales both the activation and deactivation rates of DnaA-ATP/ADP simultaneously. Increasing the switching strength shifts the transition from adder-like to non-adder behavior at high growth rates and attenuates the magnitude of the negative slope. The dashed line indicates the zero-slope reference corresponding to ideal adder behavior.

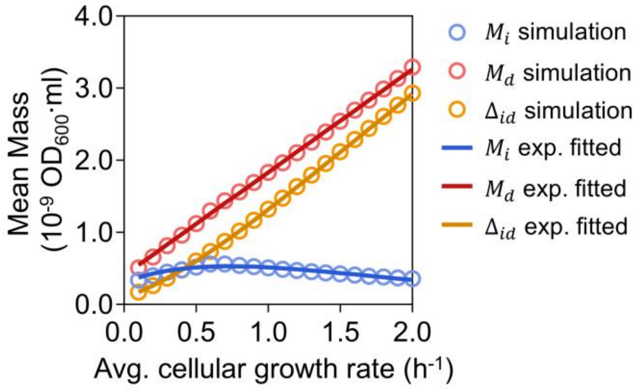


**Figure S18. Simulated initiation mass and division mass align with experimental observations.** The simulated average initiation mass ($M_i$), division mass ($M_d$) and added mass ($\Delta_{id}$) closely match the fitted curves derived from experimental data (*7*).

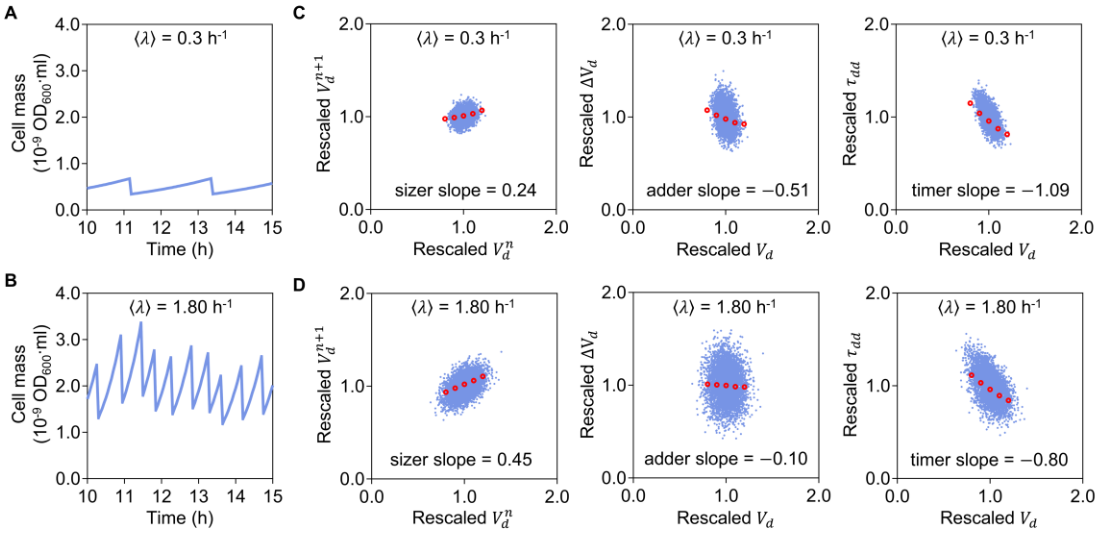


**Figure S19. Predicted cell mass dynamics and single-cell correlations between initiation and division from the IDC model.** (**AB**) Time trajectories of cell mass over 5 hours for slow or fast growth, illustrating how cell mass oscillations vary over time. (**CD**) Scatter plots for sizer, adder, and timer correlations: Sizer, rescaled division mass of the next round ($V_d^{n+1}$) versus current rescaled division mass ($V_d^n$); adder, rescaled division mass ($V_d$) versus rescaled increment during sequential division events ($\Delta V_d$); timer, rescaled increment time during sequential division events ($\tau_{dd}$) versus rescaled division mass ($V_d$). Red dots represent the average trend of the scatter plots. Sizer slope, adder slope and timer slope are obtained from linear fits of the corresponding scatter data.

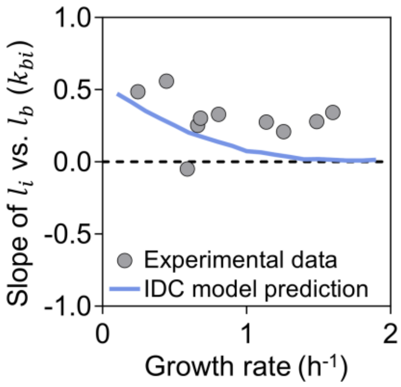


**Figure S20**. **Slope of rescaled initiation mass ($l_i$) versus rescaled birth mass ($l_b$) as a function of growth rate.** Experimental data (gray circles) show a decreasing trend in the slope with increasing growth rate. The solid blue line represents the prediction from the Initiation-to-Division Coordination (IDC) model. The dashed black line indicates zero slope, corresponding to complete independence between birth mass and initiation mass.

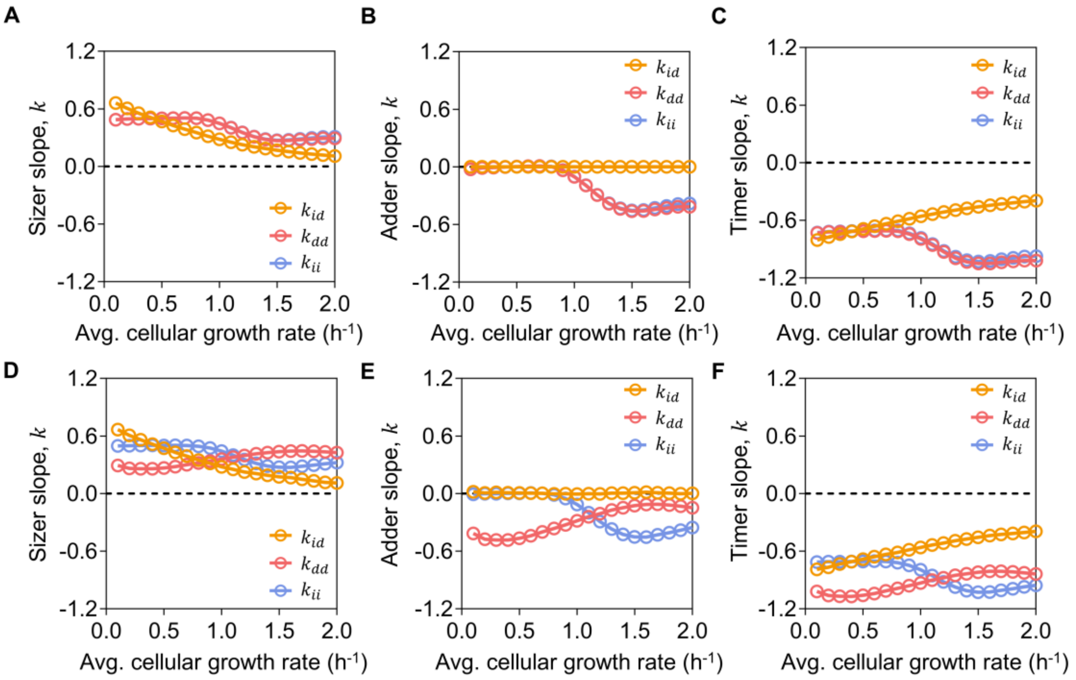


**Figure S21. Predicted growth-rate-dependent single-cell correlations from the IDC model with and without noise.** The upper row (**A-C**) shows the sizer, adder, and timer slopes as functions of growth rate without ID noise $\epsilon_{id} = 0$; whereas the lower row **(D-F)** includes growth rate fluctuations with ID noise level $\epsilon_{id} = 0.15$. Each panel compares the behavior of three key correlation parameters: $k_{ii}$, $k_{dd}$, and $k_{id}$ for single-cell correlations of three processes, initiation-to-initiation, division-to-division and initiation-to-division. This illustrates how noise affects the robustness of cell mass control mechanisms across different growth rates. The dashed line indicates the ideal sizer, adder and timer behaviors.

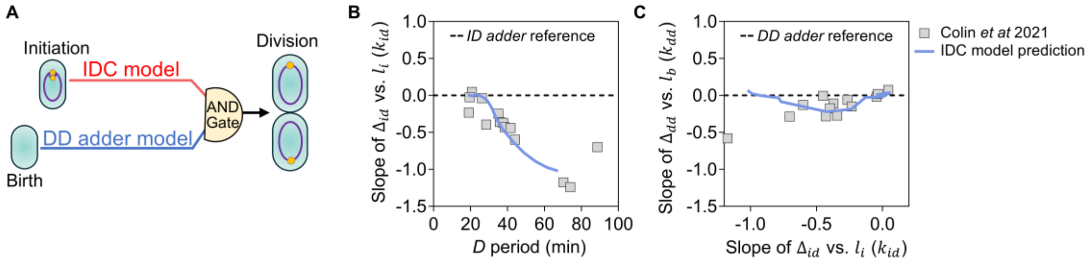


**Figure S22. Predictions of the IDC–DD concurrent model for *D*-perturbation experiments.** (**A**) Schematic of the IDC-DD concurrent model, in which cell division is jointly controlled by an ID-adder process and a DD-adder process. (**B**) ID adder slope ($k_{id}$), defined as the slope of $\Delta_{id}$ versus $l_i$, plotted as a function of the *D* period duration. (**C**) DD adder slope ($k_{dd}$), defined as the slope of $\Delta_{dd}$ versus $l_b$, plotted as a function of the ID adder slope. The blue curve represents the prediction of the IDC-DD concurrent model. Experimental data (gray squares) were extracted from Colin *et al.* (2021) (*28*).

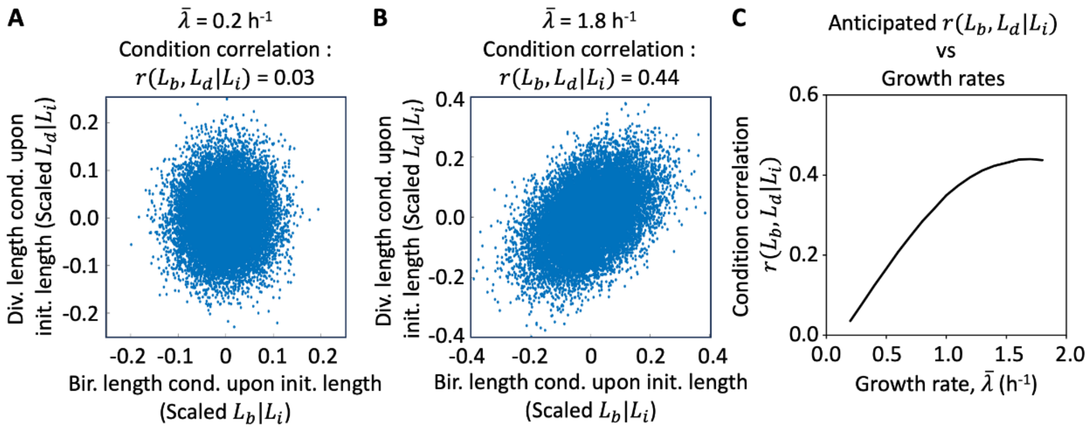


**Figure S23. Conditional independence tests linking birth, replication-initiation, and cell division within the framework of the IDC model.** (**A**) Anticipated correlation between birth mass and division mass upon fixing the initiation mass, denoted as $L_b|L_i$ and $L_d|L_i$, respectively. Data points are generated by numerical simulation of the IDC model with averaged single-cell growth rate set to 0.2 h$^{-1}$, the $r(L_b, L_d|L_i)$ represents the Pearson correlation between $L_b|L_i$ and $L_d|L_i$. (**B**), Same as panel A expect that the average single-cell growth rate ($\bar{\lambda}$) was set to 1.8 h$^{-1}$. **(C)**, The growth rate dependence of the anticipated $r(L_b, L_d|L_i)$.

**Table S1. Strains used in this study.**

| Strain | Relevant genetic marker(s) or features | Source or reference |
|---|---|---|
| MG1655 | *E.coli* K12 (AMB1655) | Liu et al.(*1*) |
| ZH101 | MG1655 Δ*fliC seqA<>seqA-mScralet* | This work |
| CLL1 | MG1655 *dnaA<> kanR intS<>*$P_{ter}$*-tetR yidA<(bla:Ptet-dnaA)>yidX* | This work |
| CLL2 | MG1655 $P_{mraZ}$ <>$P_{lac}$*-lacI* | This work |
| CLL3 | CLL1 $P_{mraZ}$ <>$P_{lac}$*-lacI* | This work |
| CLL1N | CLL1 Δ*fliC dnaN<>yPet-dnaN* | This work |
| MGAB | MG1655 Δ*thyA* Δ*deoB* | This work |

**Table S2. Reagents or resources used in this study.**

| Reagent or resource | Source | Identifier |
|---|---|---|
| Chemicals, enzymes and Critical commercial assays | | |
| DAPI | Sigma-Aldrich | Cat# D8417 |
| Rifampicin | Usbiological | Cat# R2031-70 |
| Cephalexin | Sigma-Aldrich | Cat# C4895 |
| Anhydrotetracycline (aTc) | TaKaRa | Cat# 631310 |
| Isopropyl β-D-1-thiogalactopyranoside (IPTG) | Sigma-Aldrich | Cat# I6758 |
| DpnI | NEB | Cat #R0176S |
| ClonExpress II One Step Cloning Kit | Vazyme | Cat #C112 |
| DNA Polymerase | Vazyme | Cat# P505 |
| Gel Extraction Kit | OMEGA | Cat# D2500-03 |
| Plasmid mini kit | OMEGA | Cat# D6943-03 |
| L-arabinose | Sigma-Aldrich | Cat# V900920 |
| L-Rhamnose | Sangon Biotech | Cat# A600804 |
| Thymine | Sigma-Aldrich | Cat# T0376 |
| L-Alanine | Sigma-Aldrich | Cat# A7469 |
| L-Arginine monohydrochloride | Sigma-Aldrich | Cat# V900303 |
| L-Asparagine | Sigma-Aldrich | Cat# V900458 |
| L-Aspartic acid potassium salt | Sigma-Aldrich | Cat# V900302 |
| L-Cysteine hydrochloride monohydrate | Sigma-Aldrich | Cat# C6852 |
| L-Glutamic acid monopotassium salt monohydrate | Sigma-Aldrich | Cat# 49601 |
| L-glutamine | Sigma-Aldrich | Cat# V900419 |
| Glycine | Sigma-Aldrich | Cat# G8790 |
| L-Histidine monohydrochloride monohydrate | Sigma-Aldrich | Cat# V900423 |
| L-isoleucine | Sigma-Aldrich | Cat# V900472 |
| L-leucine | Sigma-Aldrich | Cat# V900431 |
| L-Lysine monohydrochloride | Sigma-Aldrich | Cat# V900438 |
| L-Methionine | Sigma-Aldrich | Cat# V900487 |
| L-Phenylalanine | Sigma-Aldrich | Cat# V900489 |
| L-Proline | Sigma-Aldrich | Cat# V900338 |
| L-Serine | Sigma-Aldrich | Cat# V900406 |
| L-Threonine | Sigma-Aldrich | Cat# V900466 |
| L-Tryptophan | Sigma-Aldrich | Cat# V900470 |
| L-Tyrosine | Sigma-Aldrich | Cat# V900426 |
| L-Valine | Sigma-Aldrich | Cat# V900465 |
| Thiamine hydrochloride | Sigma-Aldrich | Cat# T4625 |
| Calcium pantothenate | Aladdin | Cat# C110508 |
| 4-Hydroxybenzoic acid | Sigma-Aldrich | Cat# V900794 |
| 2,3-Dihydroxybenzoic acid | Sigma-Aldrich | Cat# 126209 |
| 4-Aminobenzoic acid | Sigma-Aldrich | Cat# 100536 |
| MOPS | Sigma-Aldrich | Cat# V900306 |
| Tricine | Sigma-Aldrich | Cat# V900412 |

| | | |
|---|---|---|
| $FeSO_4 \cdot 7H_2O$ | Sigma-Aldrich | Cat# F7002 |
| $NH_4Cl$ | Sigma-Aldrich | Cat# V900222 |
| $K_2SO_4$ | Sigma-Aldrich | Cat# V900040 |
| $CaCl_2$ | Sigma-Aldrich | Cat# V900266 |
| $MgCl_2 \cdot 6H_2O$ | Sigma-Aldrich | Cat# V900020 |
| NaCl | Sigma-Aldrich | Cat# V900058 |
| Ammonium molybdate | Aladdin | Cat# A116378 |
| Boric acid | Aladdin | Cat# B111597 |
| Cobalt chloride | Aladdin | Cat# C118625 |
| Cupric sulfate | Aladdin | Cat# C110828 |
| Manganese chloride | Aladdin | Cat# M112544 |
| Zinc sulfate | Aladdin | Cat# Z165065 |
| $K_2HPO_4$ | Sigma-Aldrich | Cat# V900050 |
| Adenine | Sigma-Aldrich | Cat# V900471 |
| Cytosine | Sigma-Aldrich | Cat# V900462 |
| Guanine | Sigma-Aldrich | Cat# V900473 |
| Uracil | Sigma-Aldrich | Cat# V900439 |
| Glucose | Sigma-Aldrich | Cat# V900392 |
| Glycerol | Sigma-Aldrich | Cat# V900860 |
| Gluconate | Sigma-Aldrich | Cat# V900398 |
| Maltose | Sigma-Aldrich | Cat# V900435 |
| Sorbitol | Sigma-Aldrich | Cat# S1876 |
| Fructose | Sigma-Aldrich | Cat# F3510 |
| Succinate | Sigma-Aldrich | Cat# V900102 |
| Mannose | Aladdin | Cat# M113093 |
| Sucrose | Sigma-Aldrich | Cat# V900116 |
| Software | | |
| MATLAB | Mathworks | 2019a |
| CytExpert | Beckman | 2.4.0.28 |

**Table S3. Fitted parameters for DnaA synthesis rates ($\alpha$) over growth rates ($\lambda$).**

| | | | | | | | | | | |
|---|---|---|---|---|---|---|---|---|---|---|
| $\lambda(\mathrm{h}^{-1})$ | 0.1 | 0.2 | 0.3 | 0.4 | 0.5 | 0.6 | 0.7 | 0.8 | 0.9 | 1 |
| $\alpha(\mathrm{h}^{-1})$ | 20 | 17 | 15 | 14 | 13 | 12 | 12 | 12.43 | 12.86 | 13.29 |
| $\lambda(\mathrm{h}^{-1})$ | 1.1 | 1.2 | 1.3 | 1.4 | 1.5 | 1.6 | 1.7 | 1.8 | 1.9 | 2 |
| $\alpha(\mathrm{h}^{-1})$ | 13.72 | 14.15 | 14.58 | 15.01 | 15.44 | 15.87 | 16.3 | 16.73 | 17.16 | 17.59 |

**Table S4. Comparison between existing models.**

| Model | Initiation determination | Division determination | Reference |
|---|---|---|---|
| IDC | Initiator titration with ICC | ID adder with ICC | This study |
| IDA | II adder | DD adder | (*25*) |
| RDA | II adder | ID adder | (*26, 27*) |
| CCCP | II adder | Concurrent processes of ID timer and DD adder | (*28, 29*) |
